# Habitat and feeding ecology of a Denisovan from Late Pleistocene Taiwan

**DOI:** 10.64898/2026.08.07.743455

**Authors:** Minoru Yoneda, Chun-Hsiang Chang, Yu Itahashi, Takumi Tsutaya, Cheng-Han Sun, Cheng-Hsiu Tsai, Yousuke Kaifu

## Abstract

Denisovans, originally identified from ancient genome from Denisova Cave in Altai, were a sister group to the Neanderthals and were once widely distributed across diverse terrains in the north and south of eastern Asia^1–5^. Genomic studies suggest that there were multiple events of interbreeding between modern humans (*Homo sapiens*) and Denisovans somewhere in Asia^6^. However, little is known about Denisovan living environments, diet, ecological niche, the timing of their disappearance, and the possible coexistence with modern humans in different regions. Here we report the radiocarbon age and stable isotopic signature of Penghu 3, a large Denisovan tibia from Penghu Channel, Taiwan^7^. The results showed that Penghu 3 dates to approximately 45,000 years ago, the time when modern humans were already widespread in southern parts of Asia. This Denisovan individual inhabited a C4-dominated ecosystem, open environments such as savannahs and floodplains, or a mixture of both, and consumed a high proportion of animal protein similar to some European Neanderthals^8–10^, with no clear evidence for the use of aquatic resources. These findings have implications for the behavioral flexibility, large body size^7^, and eventual disappearance of the Denisovans.

## Main Text

Southeastern Asia (here defined as the area including Southeast, South, and southern part of East Asia) lies along the Late Pleistocene migration route of modern humans (*Homo sapiens*), who dispersed from Africa to Australia around 50,000 years ago, if not earlier^11–15^ (**Fig. 1**).

**Fig. 1.**
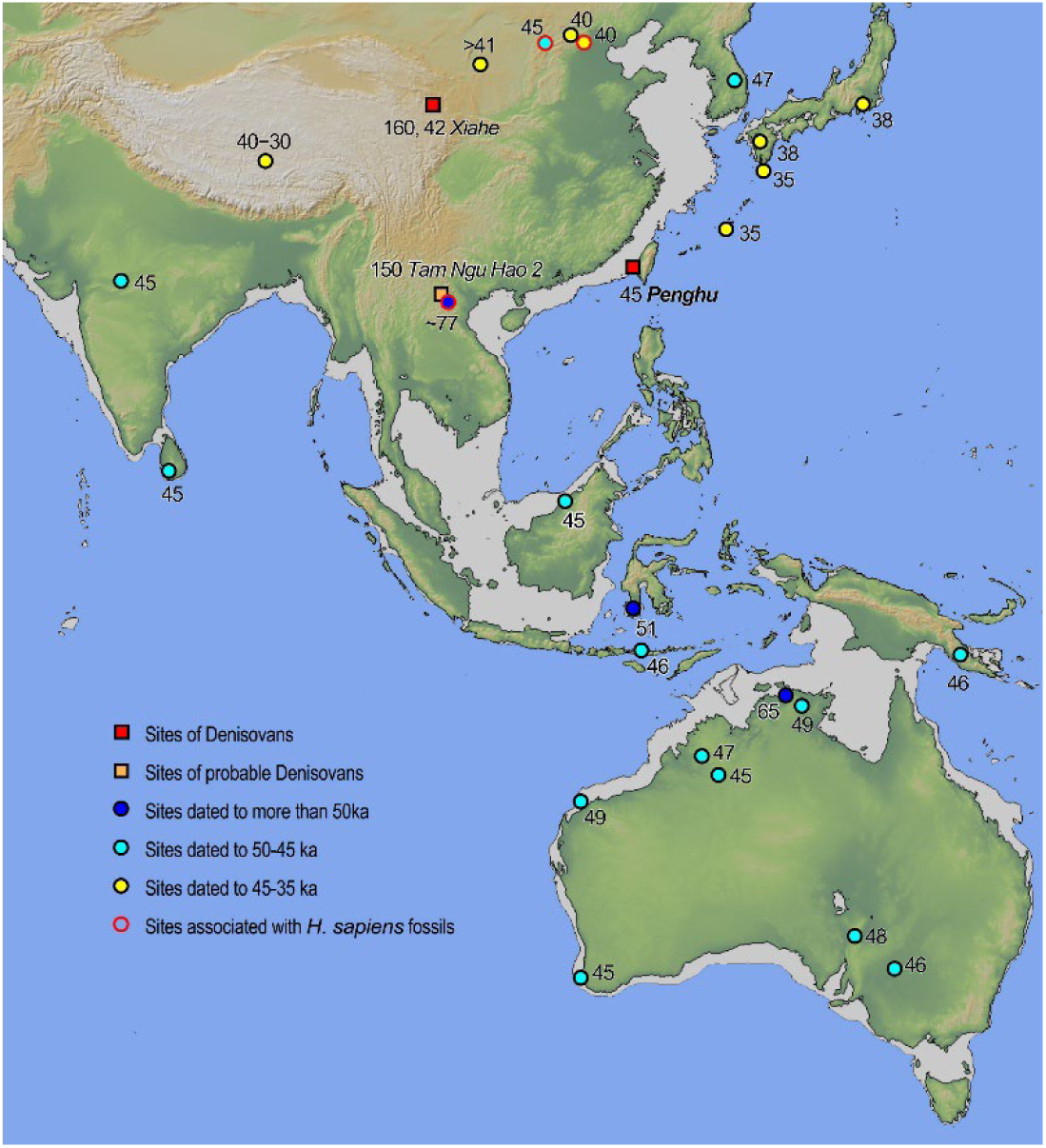
Denisovan sites around the study area and key sites presumably associated with early modern humans. Major oldest sites of modern humans in each region are shown together with their approximate ages (thousand years ago). These are Mehtakheri (India)^48^, Fa-Hien Lena (Sri Langa)^44^, Tam Pà Ling (Laos)^49^, Niah Great Cave (Borneo)^13^, Maros-Pangkep site complex (Sulawesi)^12^, Liang Bua (Flores)^14^, Ivane Vilakuav (New Guinea)^11^, Madjedbebe, Boodie, Nawarla Gabarnmang, Carpenter’s Gap, Riwi, Devils Lair, Menindeea and Warratyi (Australia)^11^, Nwya Devu (Tibet)^50^, Tianyuan Cave^51^, Xiamabei^52^, Shiyu^53^, Shuidongou^54^ (northern China), Suyanggae (South Korea)^55^, Sakitari Cave, Ishinomto and Idemaruyama (Japan)^56,57^. Site name (italic) and approximate ages for the Denisovan fossils (thousand years ago) are indicated for Denisovan sites. The base map created with GeoMapApp (www.geomapapp.org) / CC BY / CC BY (ref. ^58^).

**Fig. 2.**
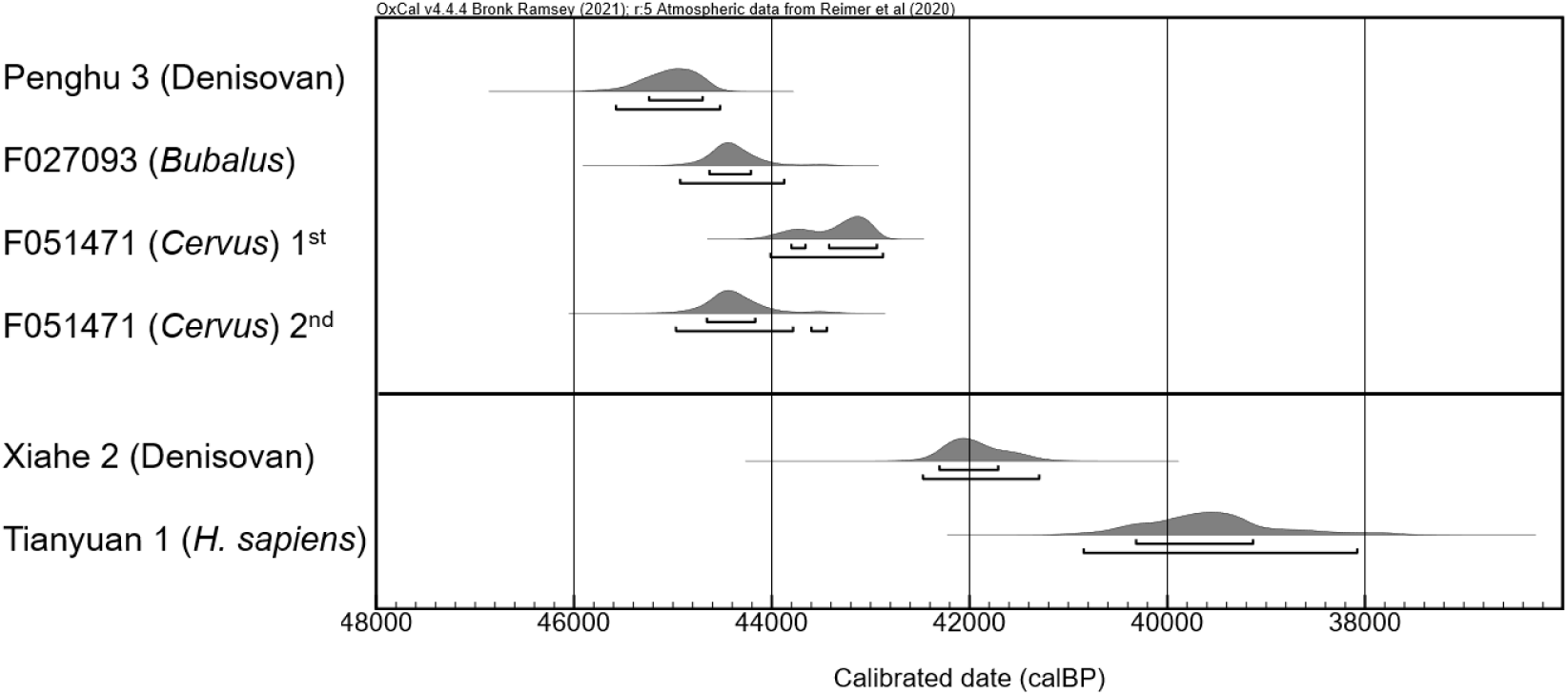
Probability distributions of calibrated radiocarbon ages on fossils from Penghu channel and other sites^38, 51^.

Genomic research suggests that this region was once inhabited by an archaic hominin group known as Denisovans^2,6^, but fossil evidence remains too limited to determine their geographic distribution or the period during which they inhabited the region^16,17^. Towards the end of the Middle Pleistocene, it is known that post-*Homo erectus* grade archaic hominin populations, represented by the fossil skulls from Maba in southern China and Narmada in India, were distributed across Southeastern Asia. However, from both molecular and morphological perspectives, it remains unclear whether these constituted a regional population of Denisovans or distinct groups of archaic hominins. Furthermore, the ages of these fossils remain unclear. The genomes of present-day Asians and Oceanians carry evidence of multiple interbreeding events with the Denisovans, one or more of which must have occurred somewhere in this region during the Late Pleistocene^2,6^. Determining the ages of Denisovan fossils is therefore essential in order to narrow down the location of such events and investigate the disappearance of the Denisovans.

Much remains unknown about the habitats and diets of ancient Asian hominins, including the Denisovans. Advances in stable isotope analysis and archaeological research in Europe have revealed that some Neanderthal populations had highly carnivorous diets^8–10^, that there was regional variation in the resource use^18^, and that they hunted medium- to large-sized animals in a manner not very different from Upper Palaeolithic modern humans. By contrast, few studies applied stable isotope analysis to human fossils in Asia, so our knowledge of the feeding ecology of archaic hominins in this region is limited^19^. Similarly, the palaeoenvironment of southeastern Asia around this time is also a subject of debate^20–24^, with important implications for regional hominin evolutionary history. For example, a recent, large-scale stable isotope study of faunal remains from this region suggests a major environmental shift that occurred around the transition from Middle to Late Pleistocene^25^. According to this study, an extensive savannah environment gave way to tropical rainforests, and this change had a direct impact on the extinction of local archaic hominins and the subsequent spread of modern humans, probably due to differences in behavioral flexibility. However, this and other scenarios^20^ remain hypothetical until more direct evidence becomes available to reconstruct the living environments of the Pleistocene hominin populations in each region.

Archaic hominin fossils and the associated faunal assemblage collected from the Penghu Channel off Taiwan (**Supplementary Note 1, Extended Data Fig. 1**, **Extended Data Table 1**) provide an opportunity to address these questions. Three hominin fossils, the Penghu 1 male mandible, the Penghu 2 femur and the Penghu 3 tibia have been recovered in this area (**Supplementary Note 2**), and palaeoproteomic analyses have revealed them to be Denisovan^4,7,26^. Based on collagen extracted from Penghu 3, this paper reports and discusses its radiocarbon (^14^C) age, palaeoenvironment and feeding ecology.

### Radiocarbon ages

We analyzed 72 faunal fossil specimens recovered from the Penghu Channel. After prescreening for nitrogen concentration using bone powder, 32 specimens that passed the criteria were selected for collagen extraction for radiocarbon (^14^C) dating and stable isotope analyses (**Supplementary Note 3**). Among these, Penghu 3 yielded the highest-quality collagen that met the criteria for acceptable preservation proposed in previous research^27,28^—namely, a bone powder nitrogen concentration of 1.7 wt%, a gelatine yield of 6.7 wt%, collagen carbon and nitrogen concentrations of 43.3 wt% and 15.6 wt%, respectively, and a collagen atomic C/N ratio of 3.2 (**Supplementary Tables 3-1and 3-2**). For the other two Denisovan specimens (Penghu 1 and Penghu 2), nitrogen was not detected in the bone powder in the prescreening (**Supplementary Table 3-1**), indicating that they retained little to no extractable collagen and were not subjected to collagen extraction.

A total of four specimens (5.6% of the prescreened materials), including Penghu 3, had atomic C/N ratios within the acceptable range (2.9–3.6), along with acceptable collagen yields and atomic concentrations, and were indicating the preservation of endogenous collagen. These specimens were selected for radiocarbon dating. Preliminary analyses of three Penghu faunal samples suggested that ultrafiltration yielded more reliable ages (**Extended Data Table 2; Extended Data Fig. 2; Supplementary Note 4**). Accordingly, radiocarbon ages were determined using ultrafiltered collagen (>30 kDa). The conventional radiocarbon age of Penghu 3 was 42,399±313 BP (TKA-30498), which was estimated to be 45,575–44,522 cal BP (95.4%) with the OxCal calibration^29^ against IntCal20^30^ (**Table 1**). Two faunal fossil specimens from Penghu produced similar ages (**Table 1**, **Fig. 3**): 41,470±297 BP (TKA-30499) for *Bubalus* (F027093), and 40,154±252 BP (TKA-30500) and 41,462±343 BP (TKA-31973) for two independent extractants of *Cervus* (F051471). A substantially older age (∼160 ka or earlier), was previously suggested for the Penghu 3 tibia based on laser ablation uranium-series dating^31^. However, the spatial distribution of uranium concentrations and ^234^U/^238^U ratios in bone is difficult to interpret in the Penghu faunal remains, because the uranium uptake/loss history in the seawater is not known^26^. It was reported that Penghu 1 and 2 have been almost entirely overprinted by a recent, pronounced influx of uranium derived from seawater^26,31^. Together with the exceptionally good collagen preservation as a Pleistocene fossil from southeastern Asia, Penghu 3 is unlikely to derive from the Middle Pleistocene and is most likely derived from Marine Isotope Stage (MIS) 3 in the Late Pleistocene, around 45 ka.

**Fig. 3.**
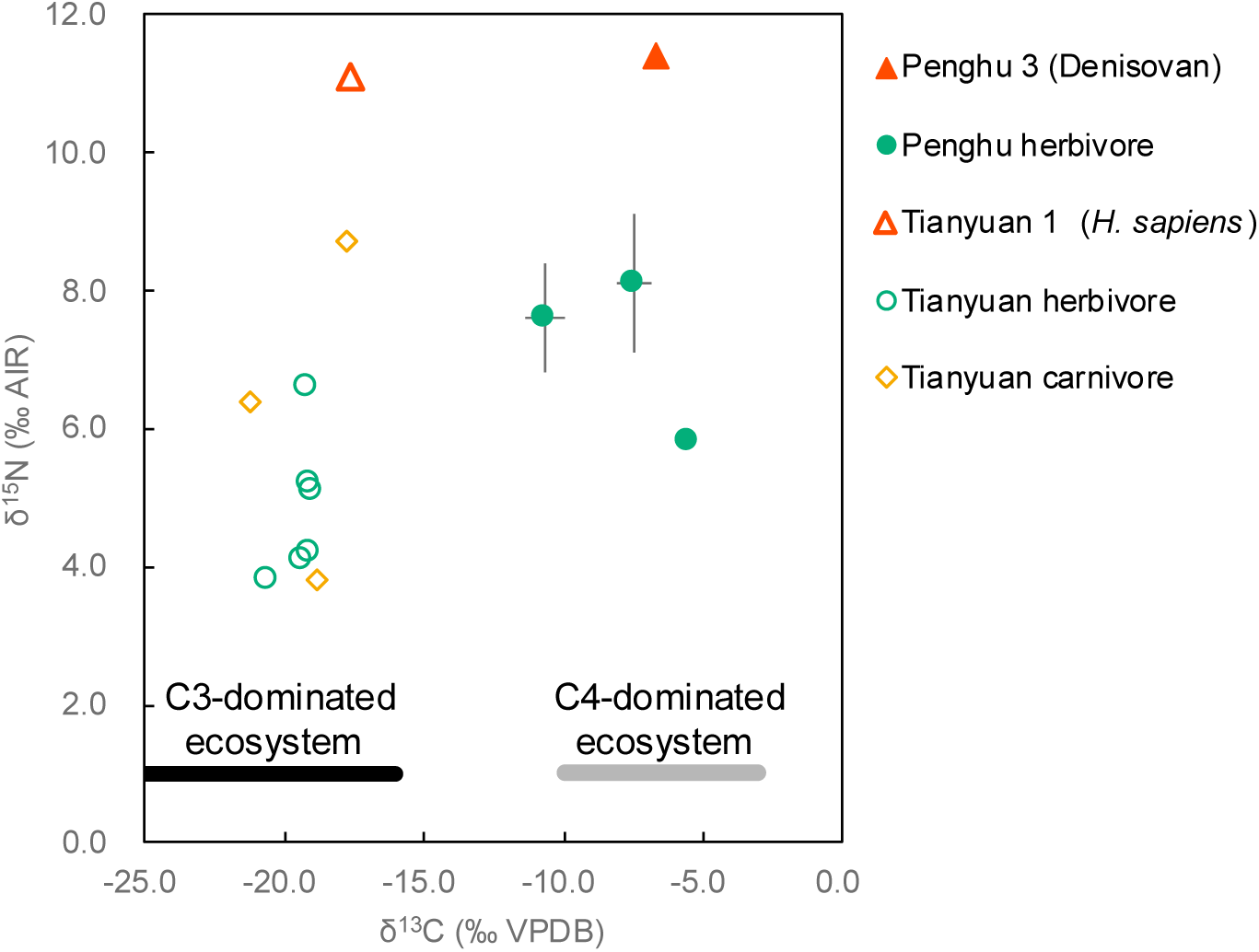
δ^13^C and δ^15^N values in bone collagen from Denisovan (Penghu 3) and faunal fossils from Penghu channel, in comparison to *Homo sapiens* and animal bones from Tianyuan Cave^43^.

**Table 1.**
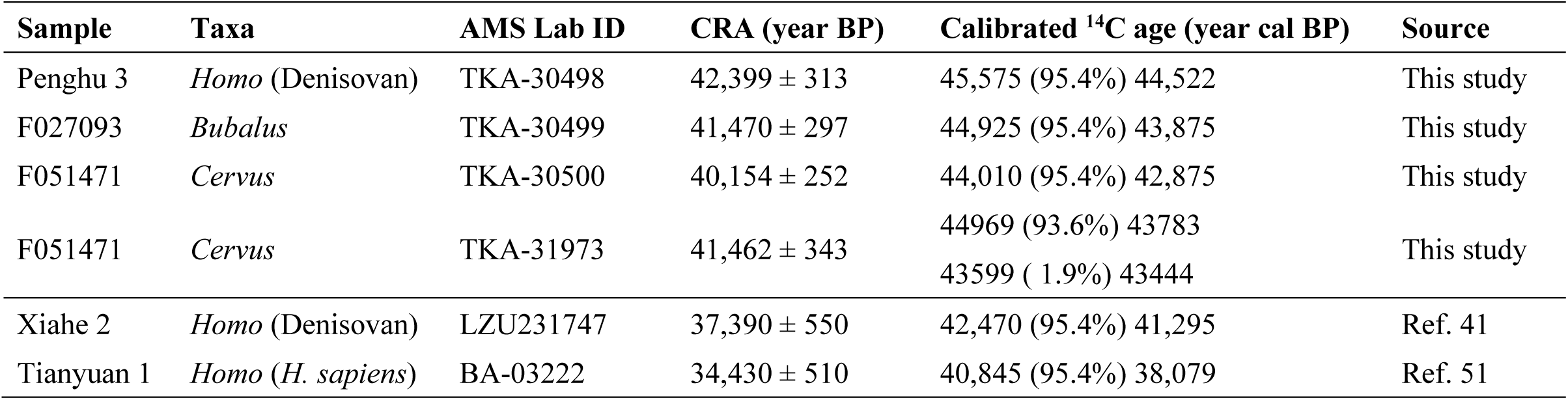
Radiocarbon ages on Penghu fossil specimens and some Asian *Homo* fossils.

| Sample | Taxa | AMS Lab ID | CRA (year BP) | Calibrated <sup>14</sup>C age (year cal BP) | Source |
| --- | --- | --- | --- | --- | --- |
| Penghu 3 | *Homo* (Denisovan) | TKA-30498 | 42,399 ± 313 | 45,575 (95.4%) 44,522 | This study |
| F027093 | *Bubalus* | TKA-30499 | 41,470 ± 297 | 44,925 (95.4%) 43,875 | This study |
| F051471 | *Cervus* | TKA-30500 | 40,154 ± 252 | 44,010 (95.4%) 42,875 | This study |
| F051471 | *Cervus* | TKA-31973 | 41,462 ± 343 | 44969 (93.6%) 43783<br>43599 ( 1.9%) 43444 | This study |
| Xiahe 2 | *Homo* (Denisovan) | LZU231747 | 37,390 ± 550 | 42,470 (95.4%) 41,295 | Ref. 41 |
| Tianyuan 1 | *Homo* (*H. sapiens*) | BA-03222 | 34,430 ± 510 | 40,845 (95.4%) 38,079 | Ref. 51 |

### Bulk stable isotope ratios

The bulk collagen carbon and nitrogen stable isotope ratios of Penghu 3 and three associated faunal specimens (two *Bubalus* and one *Cervus*), with acceptable atomic C/N ratios, are shown in **Table 2**. The bulk collagen δ^13^C values of these herbivorous mammals ranged from −5.5‰ to −10.7‰. The bulk collagen δ^13^C of Penghu 3 (−6.7‰) falls within this range (**Fig. 3**). These results indicate that Penghu 3 and the associated herbivorous mammals inhabited a C4- dominated ecosystem, representing open environments such as savannahs and floodplains, where bulk collagen δ^13^C values of approximately −3‰ to −10‰ are expected^32^. Whereas the three herbivorous mammals exhibited bulk collagen δ^15^N values of 5.5‰–8.1‰, Penghu 3 yielded a value of 11.4‰, more than 3‰ higher, indicating that this Denisovan individual occupied a relatively high trophic position (**Fig. 3**, **Supplementary Note 5**).

**Table 2.**
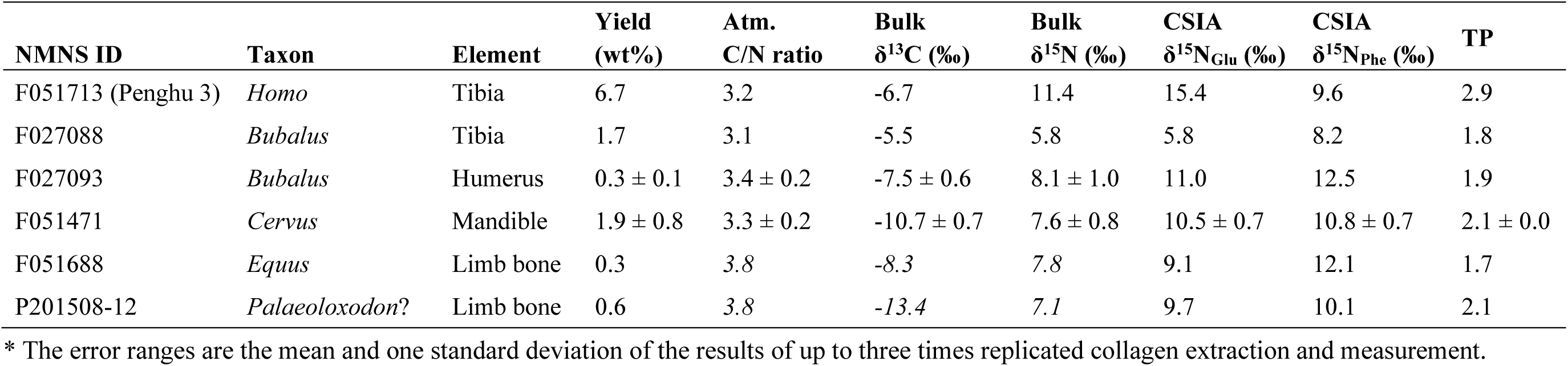
Results of preparation and isotopic analyses on Penghu fossil specimens.

### Trophic positions

In compound-specific isotope analysis (CSIA) of amino acids targeting nitrogen isotopes, trophic amino acids (e.g., Glu) exhibit substantial trophic enrichment that reflect an individual’s trophic position (TP), whereas source amino acids (e.g., Phe) show only minor ecological enrichment and retain the isotopic signature of primary producers^33^. By comparing these two groups, an animal’s TP can be estimated more accurately while avoiding the effects of baseline isotopic variation^33–35^. First, the δ^15^NPhe values, which reflect the isotopic baseline of the ecosystem, were similar between the terrestrial herbivores (8.2–12.5‰, n = 3) and Penghu 3 (9.6‰). Based on these results, Penghu 3 derived its dietary protein primarily from terrestrial animals, with no clear evidence for substantial contributions of aquatic resources, either freshwater or marine (**Fig. 4**, **Table 2**). The possibility of aquatic resource consumption cannot be excluded; however, the available data provides no support for such an interpretation. An established and validated equation^33^ that estimates the TP of terrestrial animals based on the difference between δ^15^NGlu and δ^15^NPhe (Δ^15^NGlu–Phe) was then applied. For terrestrial herbivorous mammals from Penghu (n = 3), the Δ^15^NGlu–Phe values ranged from −2.4‰ to −0.3‰ (**Table 2**). These values correspond to TP of 1.8 to 2.1, which align with the expected value (TP = 2.0) for terrestrial herbivores (**Fig. 4**). By contrast, Penghu 3 showed a Δ^15^NGlu–Phe value of 5.8‰, more than 6‰ higher than the values for the herbivores (**Table 2**). This value yielded a trophic position of 2.9 (**Table 2**). This TP value is comparable to that of terrestrial carnivores that rely exclusively on terrestrial herbivores as their nitrogen source (TP = 3.0), indicate that animal protein constituted a major component of the diet of Penghu 3. This TP value is similar to those of Neanderthals from Central Europe estimated with the same CSIA method^9,10^ (**Extended Data Table 3**). Three independent replicates of collagen extraction, amino acid derivatization, and isotopic measurement using a Penghu faunal specimen yielded standard deviations of ±0.2‰ for Δ^15^NGlu–Phe and ±0.0 for TP (**Supplementary Note 6**). These results indicate that uncertainties introduced during the experimental and mass spectrometric procedures have only negligible effects on the results.

**Fig. 4.**
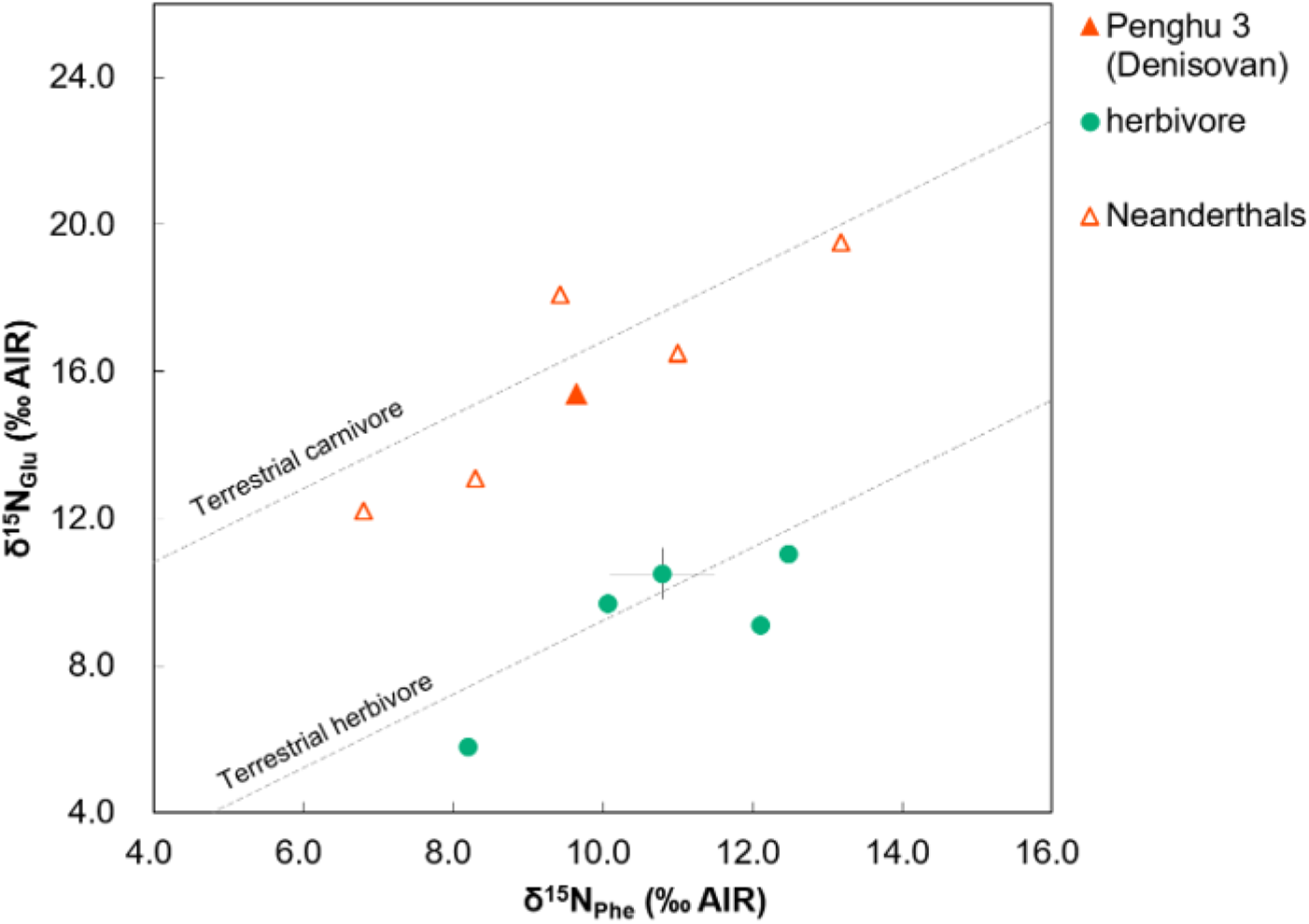
δ^15^N in phenylalanine and glutamic acids and expected TP lines for herbivore and carnivore, in comparison to European Neanderthals^9,10^.

## Discussion

Only broad age estimates, 190–130 ka or 70–10 ka, was suggested in the previous study of the Penghu 1 male Denisovan individual^26^. The radiocarbon ages reported here revealed that this faunal assemblage includes fossils of 45–43 ka. Notably, the date of 45,575–44,522 cal BP for Penghu 3 provides important evidence for understanding when the Denisovans disappeared from this region and where they interbred with modern humans.

At Denisova Cave in Altai, the youngest Denisovan fossil is genetically dated to between 52 and 76 ka^36^. At Baishiya Karst Cave on the Tibetan Plateau, a sedimentary DNA analysis implied that Denisovans used this cave until about 45 ka^37^; a Denisovan rib bone excavated from this cave (Xiahe 2) is reported to be 42 ka^38^, although this figure remains provisional due to its poor collagen preservation. If correct, these ages indicate that Denisovans survived in the two geographically distant regions, Tibet and Taiwan, until the widespread dispersal of modern humans.

In Taiwan and nearby regions, modern human sites older than 40,000 years ago have yet to be discovered. However, the known site distribution suggests that modern humans spread across a broad geographic range, from South Asia to Southeast Asia, Australia and New Guinea — between 50 and 45 ka (**Fig. 1**). Molecularly identified Denisovan fossils remain unknown from this zone, but the age of Penghu 3 does not contradict with a hypothesis that mainland southeastern Asia was one of the places where Denisovans and modern humans interbred^16^.

There is growing evidence suggesting the presence of open environments in the Pleistocene of southeastern Asia. However, its scale and how it disappeared during the Late Pleistocene, as well as the specific ecosystems used by different hominins are debated^20–25^. Our geochemical analysis of Denisovan and three other faunal fossils from the Penghu Channel revealed clear signals of a C4-dominated ecosystem. Evidence for a C4-dominated ecosystem has also been suggested based on undated elephantid specimens from Penghu^39^. Therefore, the area around the Taiwan Strait during the Late Pleistocene was characterized by open environments such as savannahs and/or the floodplains of large rivers. Some undated taxa from the Penghu assemblage, such as *Nyctereutes procyonoides*, *Ursus arctos,* and *Panthera tigris*, may be indicative of patchy or temporal existence of woodlands in this region during the Late Pleistocene (**Extended Data Table 1**, **Supplementary Note 1**). Still, the carbon isotopic signal from Penghu 3 directly suggests that the local Denisovan population primarily used an open environment.

The nitrogen isotope ratios for bulk collagen and CSIA in this study also strongly suggest that the Penghu Denisovans had a high level of meat consumption. Stable isotope and archaeological analyses have revealed that some Neanderthals consumed diets heavily reliant on animal protein and engaged in organized hunting^8–10,40^ (**Extended Data Table 3**). Consequently, some populations of the two archaic *Homo* lineages in eastern and western Eurasia similarly led a predominantly carnivorous lifestyle.

Available evidence for the feeding ecology of Asian archaic *Homo* remains limited. In Tibet, the excavated faunal remains from Baishiya Karst Cave suggest nearly full use of the local terrestrial faunal resources by Denisovans, targeting mainly on Caprinae but including megaherbivores, carnivores, and small mammals (as well as birds)^41^. In northern China, archaic *Homo* with unknown taxonomic affiliations at Xujiayao are believed to have been successful predators of large game, as evidenced by butchering marks on the bones of horses and other animals^42^.

As the burial contexts of the Penghu submarine fossils are unknown, we know little about how the Penghu Denisovans accessed animal meats. On the other hand, the morphology of the two Denisovan leg bones indicates that they were very tall individuals with estimated heights being ∼180 and ∼190 cm; the femur (Penghu 2) exhibits developed pilaster similar to Upper Palaeolithic modern human hunter-gatherers, and the tibia (Penghu 3) has a trace of healed injuries^7^. These anatomical characteristics are consistent with a highly mobile lifestyle involving regular hunting, by utilizing their powerful physiques and high activity across open terrain.

It is highly likely that a palaeo-river system existed in the Penghu Channel during the glacial period, when the area was connected to the mainland^39^. Despite this, no clear signals of aquatic resource consumption were found in Penghu 3. This contrasts with the findings from a MIS 3 *Homo sapiens* skeleton from northern China (Tianyuan 1), where high sulfur isotope ratios suggest a multifaceted subsistence strategy combining terrestrial and freshwater ecosystems^43^. Furthermore, this stands in contrast to evidence that MIS 3 *Homo sapiens* had successfully adapted to the tropical rainforest ecosystems of Sri Lanka^44^ and Borneo^45^, and utilized coastal resources in the insular environment of Wallacea^46,47^. Overall, this highlights remarkable differences in foraging flexibility between early-modern humans and the Denisovans in eastern Asia, although the data for the latter is still limited.

Some researchers hypothesized that southeastern Asian archaic *Homo* disappeared during the Late Pleistocene primarily because they were unable to flexibly adapt to expanding tropical rainforest habitats that came to dominate in this region^25^. This explanation does not contradict with our dietary and environmental reconstruction for Penghu 3. However, given the temporal overlap between southeastern Asian MIS 3 *Homo sapiens* and Denisovans demonstrated above, another explanation that needs attention is more direct competition between these hominins.

This scenario is not mutually exclusive to the environmental hypothesis, but there is little reason to assume that MIS 3 *Homo sapiens* avoided using open habitats where the Penghu Denisovans inhabited. Additional hominin fossils with secure chronometric ages and isotopic signatures would further clarify ecological difference/overlap and interaction between modern and archaic *Homo* in this region, and the disappearance of the latter.

## Methods

Transfers of Penghu 2 and 3 to Tokyo for bone sampling in 2012, 2015, 2019 and 2024 were conducted with permissions issued from the National Museum of Natural Science, Taiwan.

**Sample.** Prescreening to identify skeletal specimens with potentially well-preserved collagen was conducted on 72 Penghu fossil specimens, including three *Homo* specimens (Penghu 1, 2, and 3), from the collection of the National Museum of Natural Science (NMNS) in Taichung, Taiwan (**Supplementary Note 3**). All fossil remains were dredged from the seafloor of the Penghu Channel (Taiwan Strait), located approximately 25 km off the west coast of Taiwan, at a mean depth of 60–70 m by commercial bottom trawlers. During the colder periods with lower sea levels in the Pleistocene, this region became land that connected Taiwan to the Asian mainland. Evidence of former rivers during the Last Glacial Maximum (LGM), when global sea levels were 125–130 meters lower than today, is preserved in the area as submarine valleys with depths exceeding 100 meters.

### Radiocarbon dating

#### Prescreening

Approximately 5–10 mg of bone, dentin, and antler powder was sampled from each specimen using a tungsten carbide bit attached to a dental drill. The weight percent of carbon and nitrogen (bone C and N wt%) was subsequently measured using an elemental analyzer (Flash2000, Thermo Fisher Scientific, U.S.A.). According to a previous study, specimens with a bone nitrogen concentration (%N) of 0.76 wt% or higher and an atomic ratio of carbon to nitrogen (C/N) of 17 or lower are considered collagen samples suitable for isotope measurement^46^. However, in this study, based on preliminary investigations, a threshold of ≥0.2 bone %N was established for samples submitted for collagen extraction (**Supplementary Note 3**).

#### Collagen extraction

From the 32 samples selected through prescreening, gelatin (derived primarily from collagen) was extracted using established methods^47,48^. For several bone specimens, the gelatin extraction process was repeated multiple times, with independent extractions of gelatin samples from the same specimen (**Supplementary Note 3**). The surface deposits were physically removed by sandblasting with aluminum oxide powder and by ultrasonication in ultrapure water. The bone or dentin fragments were immersed in 1.2 mol/L HCl at 4℃ for 40 hours to demineralize the hydroxyapatite. Following a thorough rinse with ultrapure water, the fragments were immersed in a 0.1 mol/L NaOH solution for 2 hours to remove soil organic materials, such as humic and fulvic acids. After being rinsed with ultrapure water, the samples were gelatinized in a weak HCl solution (pH 4) at 90°C for 45 hours. The dissolvable gelatin was filtered through a glass filter (Whatman GF/F), and the filtered solution was freeze-dried to obtain samples for isotopic measurements.

#### Ultrafiltration

For the four successfully extracted gelatin samples, including Penghu 3, with the indicators of well-preserved collagen, a subsequent purification process via ultrafiltration (UF) was employed prior to radiocarbon dating. The use of UF is particularly effective for removing exogenous organic matter trapped within the collagen fibers and for purifying collagen for older bones/dentins older than 40,000 years^49^. In Penghu fauna, Vivaspin 20 30 kDa MWCO (GH Healthcare) successfully removed exogenous carbon compounds and yielded more precise ^14^C ages derived from endogenous collagen (**Extended Data Fig. 2; Extended Data Table 2; Supplementary Note 4**). Considering the possible contamination of carbon atoms from residual humectant on the membrane, the ultrafilter was thoroughly washed before use.

First, a sufficient volume of ultrapure water was passed through the membrane^50^. Subsequently, the membrane unit was ultrasonicated in ultrapure water for 1 h, and this procedure was repeated twice. No traces of carbon were detected with an elemental analyzer in the water that passed through the ultrafiltration membrane after this cleaning process.

#### Accelerator mass spectrometry (AMS) dating

The extracted gelatin samples were graphitized using two compatible methods, and the ^14^C abundance was measured by accelerator mass spectrometry (AMS) to determine the conventional radiocarbon age (CRA) calculated in units of before present (BP)^51^. In one graphitization method, the gelatin was enclosed in a quartz glass tube along with copper oxide and Sulfix. The tube was heated at 850°C for 3 h, generating a gas comprising carbon dioxide^52^. The combustion gases from each sample were introduced into a vacuum line, and CO2 was purified. Subsequently, the purified CO2 was mixed with hydrogen and iron powder catalyst, sealed in a reaction vessel, and heated at 650°C for 6 h, to obtain graphite^53^. In another graphitization method, gelatin was introduced into an elemental analyzer (vario ISOTOPE select, Elementar Analysensysteme GmbH, Germany) to perform combustion and CO2 purification. Graphite was then obtained through purification via a vacuum line and reduction with hydrogen and an iron catalyst^54^.

The Compact-AMS (National Electrostatics Corp., U.S.A.) at the Laboratory of Radiocarbon Dating, The University Museum, The University of Tokyo (UMUT), was used to measure five standard and three background targets concurrently with the unknown samples in a typical measurement. The standard material used for correction was HOxII, distributed by NIST. The background was established using a commercial oxalic acid (Wako-OX) as a working standard, which is characterized by ^14^C content lower than that of graphite derived from the standard material marble (IAEA-C1) (**Extended Data Fig. 3**). A probability distribution of the calibrated date was estimated using the measured CRA with the IntCal20^30^ and OxCal 4.4 software^29^.

### Stable isotope analyses

#### Carbon and nitrogen isotope analysis of bulk collagen

The stable carbon and nitrogen isotope ratios, as well as weight percentages of carbon and nitrogen, were measured at UMUT. Approximately 0.4 mg of extracted gelatine were measured by an EA-IRMS (Flash2000 elemental analyzer and Delta V Advantage isotope ratio mass spectrometer, Thermo Fisher Scientific Inc., U.S.A.). The nitrogen isotope ratios (δ^15^N) were normalized to the value of the standard atmosphere (AIR), and the carbon isotope ratios (δ^13^C) were normalized to the value of the Vienna Peedee Belemnite (VPDB). The measurement errors of the instrument were determined to be < ±0.1 ‰ for δ^15^Ncol and < ±0.1 ‰ for δ^13^Ccol in one standard deviation (SD) by repeated measurement of working amino acid standards (alanine, glycine, and histidine, provided by SI Science Inc., Japan) with known values (**Supplementary Note 5**).

Because of prolonged turnover rates, collagen isotope ratios of cortical limb bones reflect the average of an individual’s diet over a long period, approximately spanning the past 10–20 years^55^. The consumption of C4 plants—which are adapted to open, dry habitats—or marine products leads to heavier δ^13^C values^56^. Abundance of heavy nitrogen isotopes is greater in the higher positions of the food chain, and higher consumers in long food chains in aquatic ecosystems tend to have higher δ^15^N values than terrestrial animals with shorter food chains^57^.

#### Compound-specific isotope analysis (CSIA) of amino acids

In this study, collagen was derivatized using a modified method^58^ based on the methodology of ref. ^34^ for isolation by gas chromatography and nitrogen isotope analysis. Initially, 2 mg of collagen was hydrolyzed in 12 mol/L HCl at 110°C for 12 h. Then, after lipid removal with hexane and dichloromethane (3/2, v/v) wash, the solution was dried under a continuous N2 flow. Subsequently, thionyl chloride and 2-propanol (1/4, v/v) were added, and the samples were heated at 110°C for 2 h. After heating, the samples were dried again and then heated at 110°C for 2 h with pivaloyl chloride and dichloromethane (1/4, v/v) to derivatize amino acids to highly volatile compounds. The solution of derivatized amino acids, recovered using a hexane and dichloromethane mixture (3/2, v/v), was adjusted to the appropriate concentration for isotopic measurement. The nitrogen isotope ratios were subsequently measured using a GC-C-IRMS (TRACE1310GC, coupled to a Thermo Finnigan Delta V Advantage IRMS via a GC Isolink II interface, Thermo Fisher Scientific Inc., USA) at UMUT. The measurement error (1σ) of the instrument, evaluated using nine amino acid standards with guaranteed values (SI Science Co., Ltd., Japan; **Supplementary Note 6**), was ±0.3‰ to ±0.9‰.

## Acknowledgments

C-H.C. sincerely thanks Mr. Li-Ren Hou for his generous donation of the Penghu faunal fossils to the National Museum of Natural Science, Taiwan. We also thank Takashi Gakuhari, Masahiro Ozaki, Takayuki Omori, Tadao Kanesawa, and Kohei Yamazaki for sampling and isotopic analysis, and Katherine Hampson for editorial assistance. This work was supported by JSPS KAKENHI Grant Numbers 15H05969, 22101005 and 22H00421, JST FOREST program Grant Number JPMJFR233D, NSTC Grant Number 112-2116-M-178-001-, and NTU FD107028.

## Author contributions

M.Y, C.-H.C. and Y.K. conceived the study. M.Y, Y.I. collected isotopic data. M.Y. and Y.K. wrote the main text manuscript with critical input from the remaining authors. M.Y., C.-H.C., Y.I., T.T., C.-H.S., C.-H.T. and Y.K. wrote the SI manuscript.

---

**Extended Data Fig. 1.**
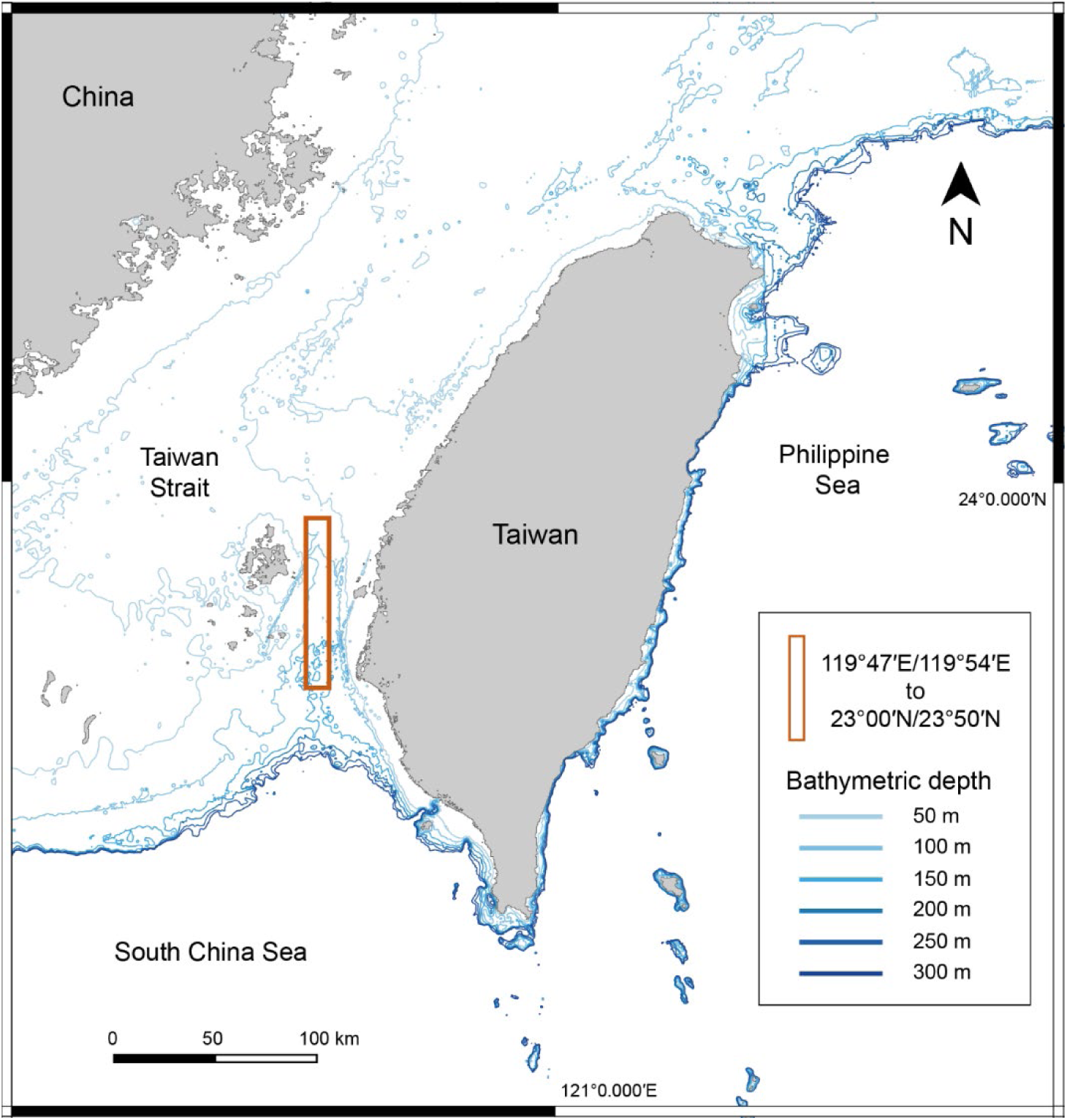
Submarine topography around the Penghu Channel. The red square indicates the main area of fossil accumulation on the sea floor.

**Extended Data Fig. 2.**
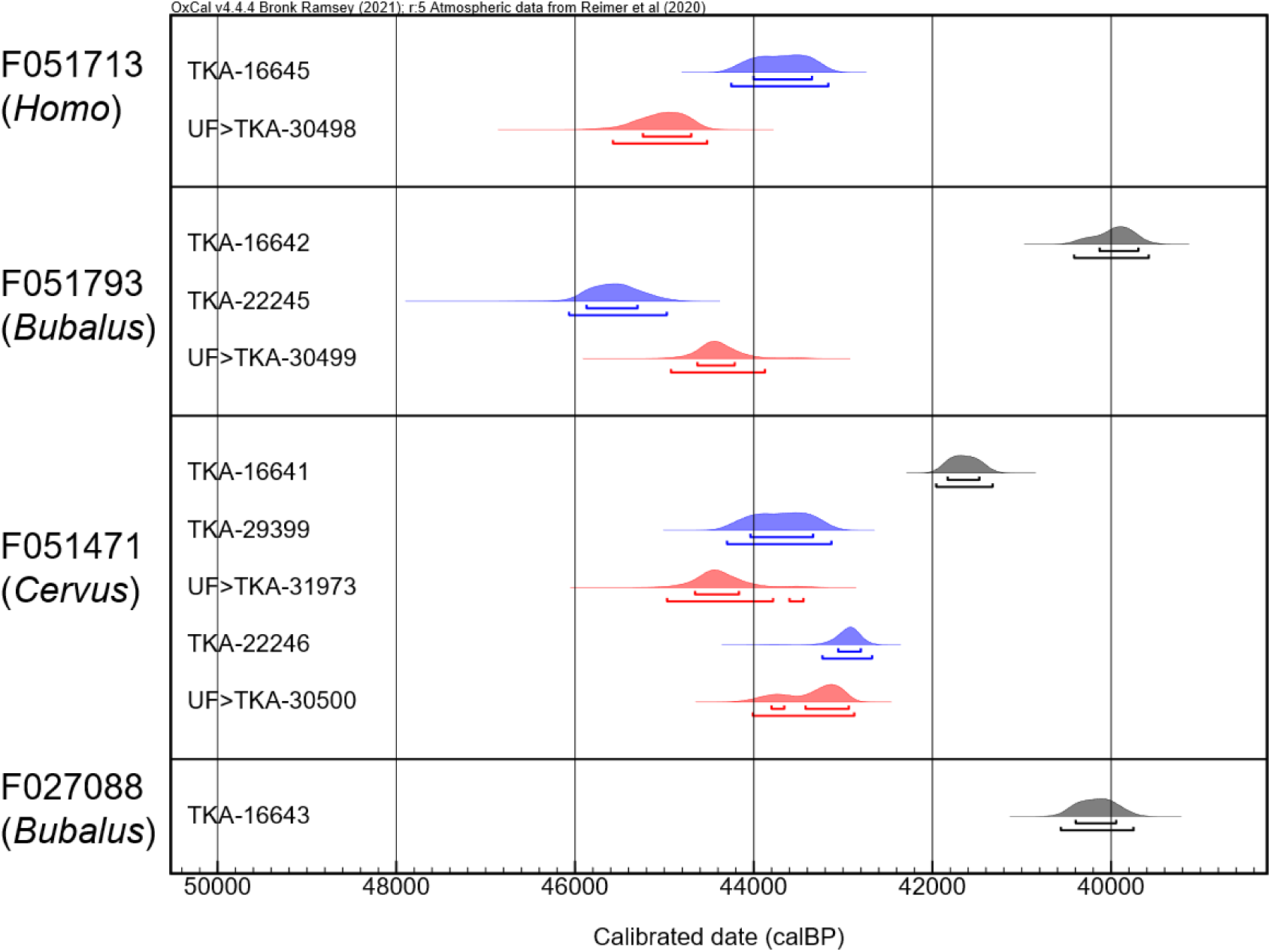
Calibrated radiocarbon ages of Penghu fossils. Calibrated ^14^C ages before (blue) and after (red) ultrafiltration, together with those of other extractants without ultrafiltration from the same specimen (gray). Up to three extractants were obtained from each specimen. The identifiers starting with “F” represent specimens. The identifiers beginning with “TKA” represent the AMS laboratory identifiers, and “UF” means ultrafiltration. The convergence of calibrated ^14^C ages among extractants from different specimens following ultrafiltration suggests that this step successfully removed exogenous carbon compounds and yielded more accurate ages derived from endogenous collagen.

**Extended Data Fig. 3.**
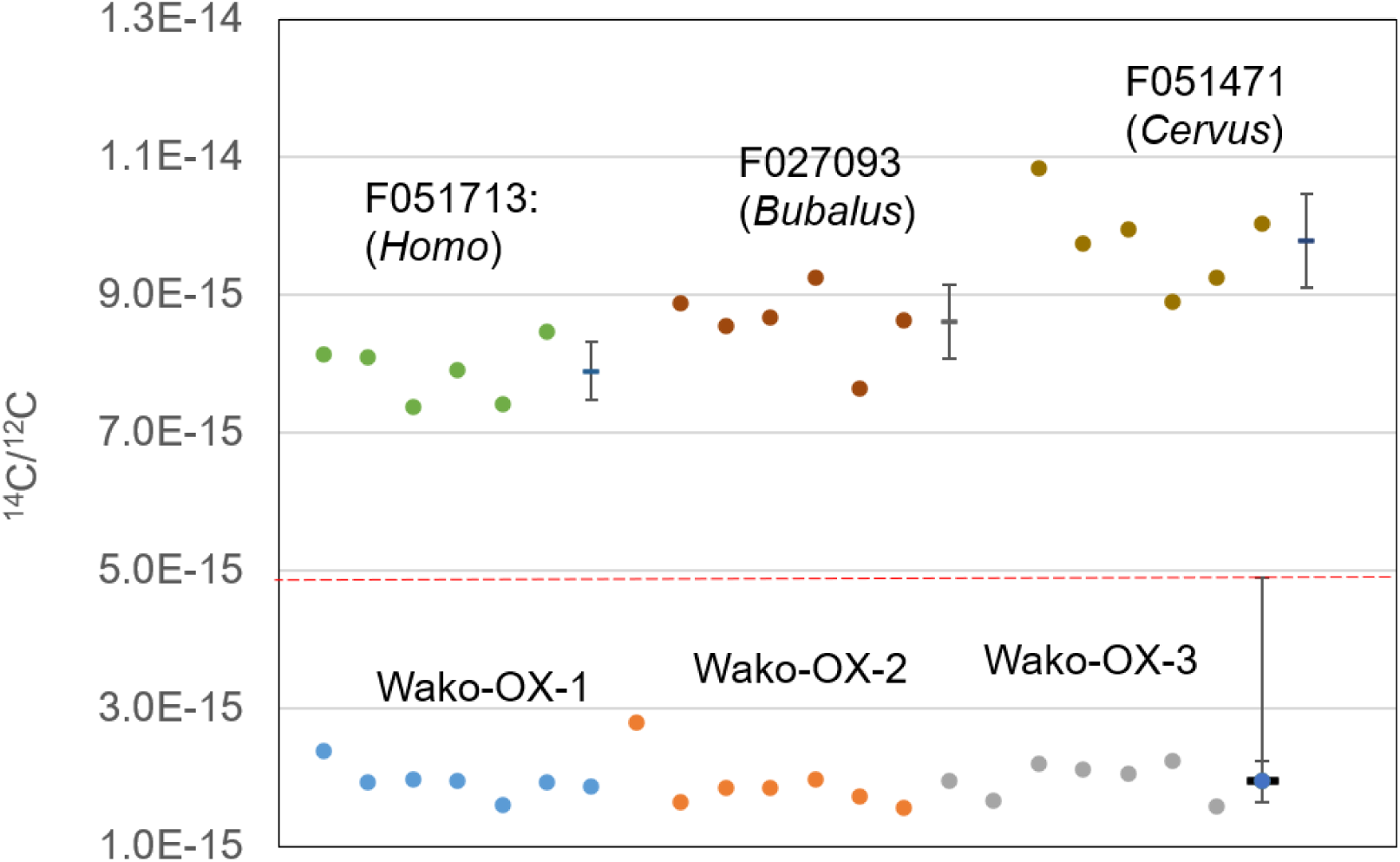
Radiocarbon contents of the background and Penghu fossil sample. Measured ^14^C contents of the background (Wako-OX) and three Penghu fossil samples (F051713, F027093, and F051471) are shown. The dots at the right-hand end of each sample plot indicate the mean value and 1 SD error bars. For Wako-OX, 10 SD error bar and a dashed red line indicate the limit of quantification. Although the ^14^C contents of the Penghu fossils lie close to the quantification limit, their ^14^C concentrations were measured reliably in the AMS system.

**Extended Data Table 1.**
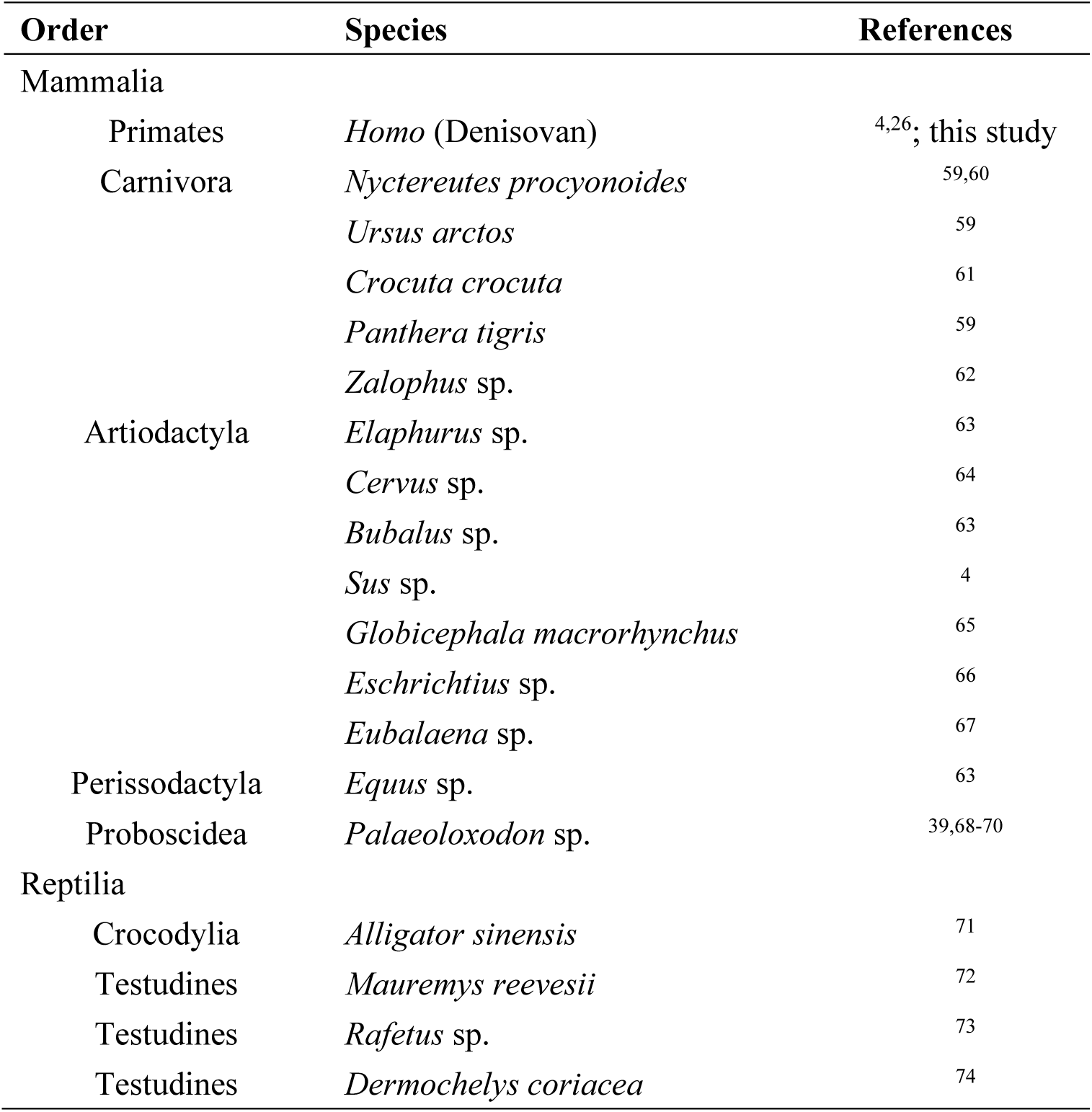
List of major fossil vertebrates from the Penghu assemblage. Given the taxonomic uncertainty of various fossil materials and ongoing studies, some species are listed as unspecified, indicated by ‘sp.’ after the genus name.

**Extended Data Table 2.**
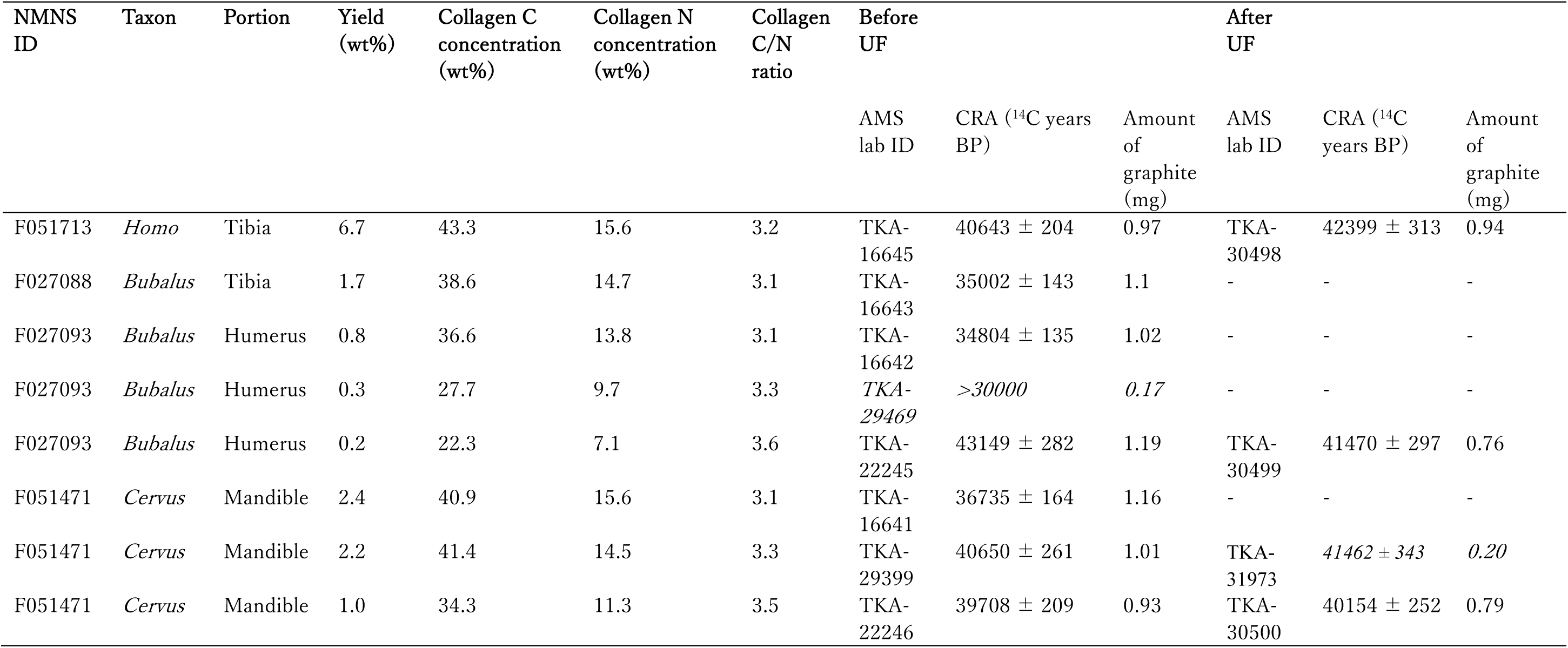
Bulk stable isotopic and radiocarbon results of the Penghu fossils with the acceptable collagen extraction measures. Results of the same extractants before and after applying ultrafiltration (UF) are shown for some samples.

| NMNS ID | Taxon | Portion | Yield (wt%) | Collagen C concentration (wt%) | Collagen N concentration (wt%) | Collagen C/N ratio | Before UF |  |  | After UF |  |  |
| --- | --- | --- | --- | --- | --- | --- | --- | --- | --- | --- | --- | --- |
|  |  |  |  |  |  |  | AMS lab ID | CRA (<sup>14</sup>C years BP) | Amount of graphite (mg) | AMS lab ID | CRA (<sup>14</sup>C years BP) | Amount of graphite (mg) |
| F051713 | *Homo* | Tibia | 6.7 | 43.3 | 15.6 | 3.2 | TKA-16645 | 40643 ± 204 | 0.97 | TKA-30498 | 42399 ± 313 | 0.94 |
| F027088 | *Bubalus* | Tibia | 1.7 | 38.6 | 14.7 | 3.1 | TKA-16643 | 35002 ± 143 | 1.1 | - | - | - |
| F027093 | *Bubalus* | Humerus | 0.8 | 36.6 | 13.8 | 3.1 | TKA-16642 | 34804 ± 135 | 1.02 | - | - | - |
| F027093 | *Bubalus* | Humerus | 0.3 | 27.7 | 9.7 | 3.3 | TKA-29469 | >30000 | 0.17 | - | - | - |
| F027093 | *Bubalus* | Humerus | 0.2 | 22.3 | 7.1 | 3.6 | TKA-22245 | 43149 ± 282 | 1.19 | TKA-30499 | 41470 ± 297 | 0.76 |
| F051471 | *Cervus* | Mandible | 2.4 | 40.9 | 15.6 | 3.1 | TKA-16641 | 36735 ± 164 | 1.16 | - | - | - |
| F051471 | *Cervus* | Mandible | 2.2 | 41.4 | 14.5 | 3.3 | TKA-29399 | 40650 ± 261 | 1.01 | TKA-31973 | 41462 ± 343 | 0.20 |
| F051471 | *Cervus* | Mandible | 1.0 | 34.3 | 11.3 | 3.5 | TKA-22246 | 39708 ± 209 | 0.93 | TKA-30500 | 40154 ± 252 | 0.79 |

**Extended Data Table 3.**
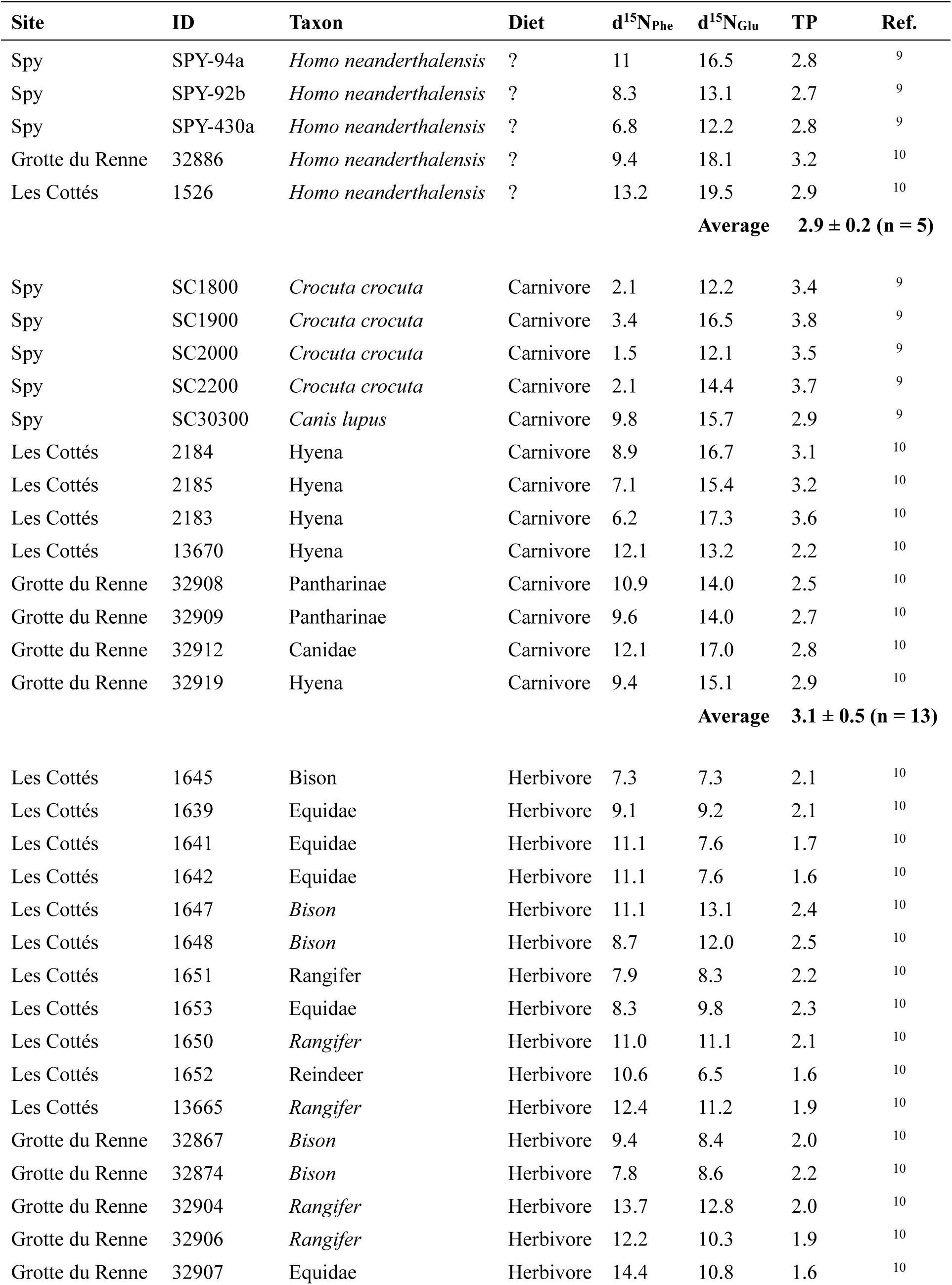
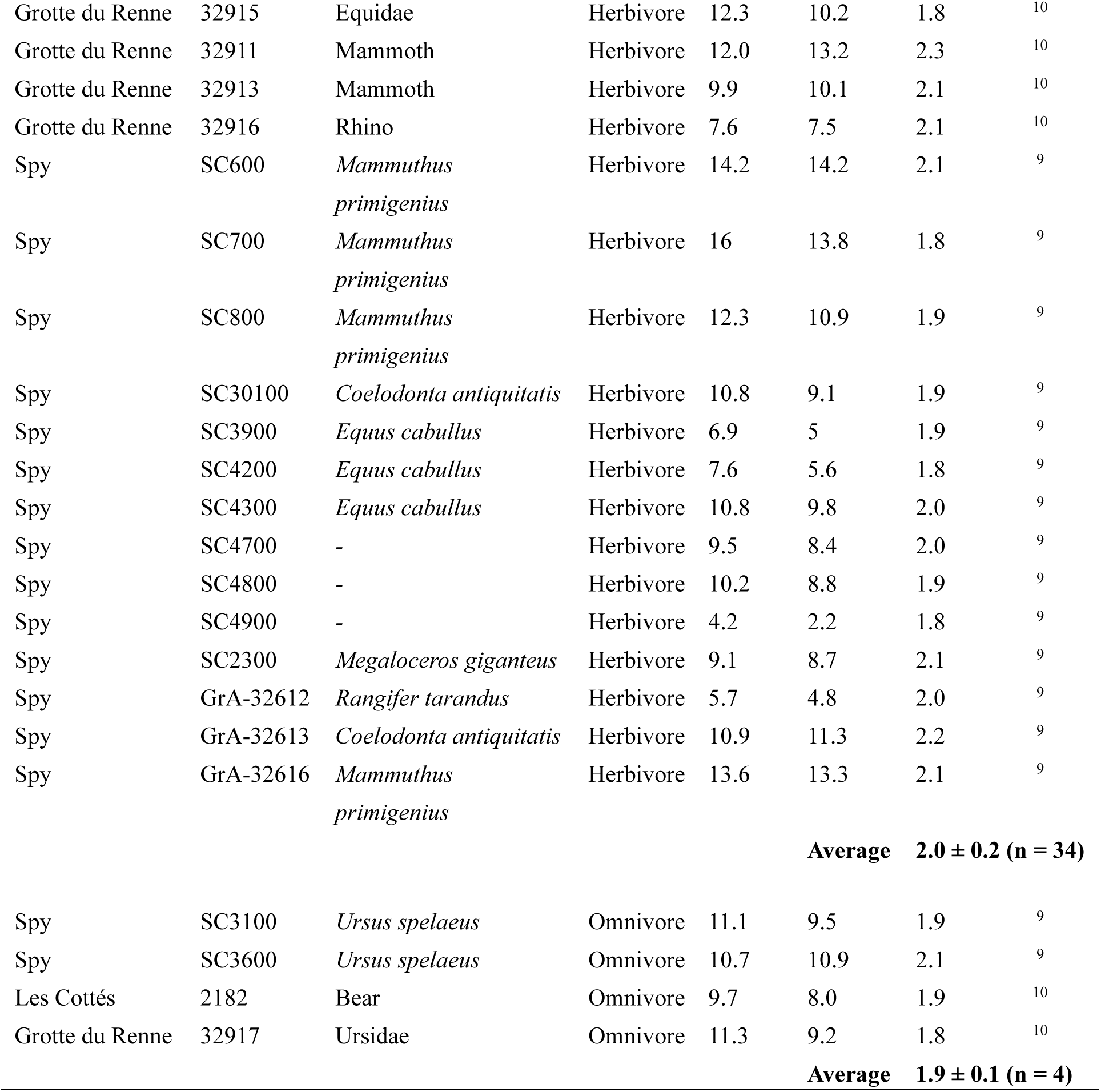
CSIA results and calculated TPs of previously reported Neanderthal individuals and associated fauna. The average TP and its 1 SD ranges are shown for each category of the animal diet.

## References

1 Reich, D. et al. Genetic history of an archaic hominin group from Denisova Cave in Siberia. Nature 468, 1053–1060 (2010).

2 Peyregne, S., Slon, V. & Kelso, J. More than a decade of genetic research on the Denisovans. Nature Reviews Genetics 25, 83–103 (2024).

3 Chen, F. et al. A late Middle Pleistocene Denisovan mandible from the Tibetan Plateau. Nature 569, 409–412 (2019).

4 Tsutaya, T. et al. A male Denisovan mandible from Pleistocene Taiwan. Science 388, 176–180 (2025).

5 Fu, Q. et al. The proteome of the late Middle Pleistocene Harbin individual. Science 389, 704–707 (2025).

6 Jacobs, G. S. et al. Multiple Deeply Divergent Denisovan Ancestries in Papuans. Cell 177, 1010–1021 e1032 (2019).

7. Kaifu, Y., et al. Denisovan leg bones from Taiwan reveal large body size. bioRxib (2026).

8 Richards, M. P. & Trinkaus, E. Isotopic evidence for the diets of European Neanderthals and early modern humans. Proc Natl Acad Sci U S A 106, 16034–16039 (2009).

9 Naito, Y. I. et al. Ecological niche of Neanderthals from Spy Cave revealed by nitrogen isotopes of individual amino acids in collagen. J Hum Evol 93, 82–90 (2016).

10 Jaouen, K. et al. Exceptionally high delta(15)N values in collagen single amino acids confirm Neandertals as high-trophic level carnivores. Proc Natl Acad Sci U S A 116, 4928–4933 (2019).

11 O’Connell, J. F. et al. When did Homo sapiens first reach Southeast Asia and Sahul? Proc Natl Acad Sci USA 115, 8482–8490 (2018).

12 Oktaviana, A. A. et al. Narrative cave art in Indonesia by 51,200 years ago. Nature 631, 814–818 (2024).

13 Higham, T. F. G. et al. Radiocarbon dating of charcoal from tropical sequences: results from the Niah Great Cave, Sarawak, and their broader implications. Journal of Quaternary Science 24, 189–197 (2009).

14 Sutikna, T. et al. The spatio-temporal distribution of archaeological and faunal finds at Liang Bua (Flores, Indonesia) in light of the revised chronology for Homo floresiensis. J Hum Evol 124, 52–74 (2018).

15. Kaifu, Y., et al. Modern human teeth unearthed from below the approximately 128,000-year-old level at Punung, Java: A case highlighting the problem of recent intrusion in cave sediments. J Hum Evol 163, 103122 (2022).

16 Kaifu, Y. & Athreya, S. Diversity and evolution of archaic eastern Asian hominins: A synthetic model of the fossil and genetic records. PaleoAnthropology 2025, 370−397 (2025).

17 Sawafuji, R., Tsutaya, T., Takahata, N., Pedersen, M. W. & Ishida, H. East and Southeast Asian hominin dispersal and evolution: A review. Quaternary Science Reviews 333 (2024).

18 Weyrich, L. S. et al. Neanderthal behaviour, diet, and disease inferred from ancient DNA in dental calculus. Nature 544, 357–361 (2017).

19 Kubat, J. et al. Dietary strategies of Pleistocene Pongo sp. and Homo erectus on Java (Indonesia). Nat Ecol Evol 7, 279–289 (2023).

20 Bird, M. I., Taylor, D. & Hunt, C. Palaeoenvironments of insular Southeast Asia during the Last Glacial Period: a savanna corridor in Sundaland? Quaternary Science Reviews 24, 2228–2242 (2005).

21 Cannon, C. H., Morley, R. J. & Bush, A. B. The current refugial rainforests of Sundaland are unrepresentative of their biogeographic past and highly vulnerable to disturbance. Proc Natl Acad Sci U S A 106, 11188–11193 (2009).

22 Morley, R. J. in Biotic Evolution and Environmental Change in Southeast Asia Ch. 4, 79–114 (2012).

23 McGrath, S. M., Clemens, S. C. & Huang, Y. Pleistocene Sunda Shelf submersion-exposure cycles initiate vegetation Walker Circulation feedback. Geology 51, 1053–1056 (2023).

24 Hamilton, R. et al. Forest mosaics, not savanna corridors, dominated in Southeast Asia during the Last Glacial Maximum. Proc Natl Acad Sci U S A 121, e2311280120 (2024).

25 Louys, J. & Roberts, P. Environmental drivers of megafauna and hominin extinction in Southeast Asia. Nature 586, 402–406 (2020).

26 Chang, C. H. et al. The first archaic Homo from Taiwan. Nat. Commun. 6, 6037 (2015).

27 DeNiro, M. J. Postmortem preservation and alteration of in vivo bone collagen isotope ratios in relation to palaeodietary reconstruction. Nature 317, 806–809 (1985).

28 van Klinken, G. J. Bone Collagen Quality Indicators for Palaeodietary and Radiocarbon Measurements. Journal of Archaeological Science 26, 687–695 (1999).

29 Bronk Ramsey, C. Radiocarbon Calibration and Analysis of Stratigraphy: The OxCal Program. Radiocarbon 37, 425–430 (1995).

30 Reimer, P. J. et al. The IntCal20 Northern Hemisphere Radiocarbon Age Calibration Curve (0–55 cal kBP). Radiocarbon 62, 725–757 (2020).

31 Grün, R. & Stringer, C. Direct dating of human fossils and the ever-changing story of human evolution. Quaternary Science Reviews 322 (2023).

32 van der Merwe, N. J. & Vogel, J. C. 13C content of human collagen as a measure of prehistoric diet in woodland North America. Nature 276, 815–816 (1978).

33 Ohkouchi, N. et al. Advances in the application of amino acid nitrogen isotopic analysis in ecological and biogeochemical studies. Organic Geochemistry 113, 150–174 (2017).

34. Chikaraishi, Y., Ogawa, N. O. & Ohkouchi, N. in Earth, life, and isotopes (eds N. Ohkouch, I. Tayasu, & K. Koba) 37–51 (Kyoto University Press, 2010).

35 Chikaraishi, Y., Ogawa, N. O., Doi, H. & Ohkouchi, N. 15N/14N ratios of amino acids as a tool for studying terrestrial food webs: a case study of terrestrial insects (bees, wasps, and hornets). Ecological Research 26, 835–844 (2011).

36 Douka, K. et al. Age estimates for hominin fossils and the onset of the Upper Palaeolithic at Denisova Cave. Nature 565, 640–644 (2019).

37 Zhang, D. et al. Denisovan DNA in Late Pleistocene sediments from Baishiya Karst Cave on the Tibetan Plateau. Science 370, 584–587 (2020).

38 Xia, H., Li, Y., Zhang, D. & Chen, F. New insights from the latest Denisovan fossil discovery on the Tibetan Plateau. Chinese Science Bulletin 69, 5155–5160 (2024).

39 Biswas, D. S., Banerjee, Y., Baker, D., Chang, C. H. & Tsai, C. H. A glimpse into a vanished ecosystem: reconstructing diet and palaeoenvironment of Palaeoloxodon from the Pleistocene of Taiwan. R Soc Open Sci 12, 250935 (2025).

40 Hutson, J. M. et al. Revised age for Schoningen hunting spears indicates intensification of Neanderthal cooperative behavior around 200,000 years ago. Sci Adv 11, eadv0752 (2025).

41 Xia, H. et al. Middle and Late Pleistocene Denisovan subsistence at Baishiya Karst Cave. Nature 632, 108–113 (2024).

42 Norton, C. J. & Gao, X. Hominin-carnivore interactions during the Chinese Early Paleolithic: taphonomic perspectives from Xujiayao. J Hum Evol 55, 164–178 (2008).

43 Hu, Y. et al. Stable isotope dietary analysis of the Tianyuan 1 early modern human. Proc Natl Acad Sci U S A 106, 10971–10974 (2009).

44 Wedage, O. et al. Specialized rainforest hunting by Homo sapiens ∼45,000 years ago. Nature communications 10, 739 (2019).

45 Barker, G. et al. The ‘human revolution’ in lowland tropical Southeast Asia: the antiquity and behavior of anatomically modern humans at Niah Cave (Sarawak, Borneo). J Hum Evol 52, 243–261 (2007).

46 O’Connor, S., Ono, R. & Clarkson, C. Pelagic fishing at 42,000 years before the present and the maritime skills of modern humans. Science 334, 1117–1121 (2011).

47 Roberts, P. et al. Isotopic evidence for initial coastal colonization and subsequent diversification in the human occupation of Wallacea. Nature communications 11, 2068 (2020).

48 Mishra, S., Chauhan, N. & Singhvi, A. K. Continuity of microblade technology in the Indian subcontinent since 45 ka: Implications for the dispersal of modern humans. PLoS ONE 8(**7**): e69280 (2013).

49 Freidline, S. E. et al. Early presence of Homo sapiens in Southeast Asia by 86-68 kyr at Tam Pa Ling, Northern Laos. Nature communications 14, 3193 (2023).

50 Zhang, X. L. et al. The earliest human occupation of the high-altitude Tibetan Plateau 40 thousand to 30 thousand years ago. Science 362, 1049–1051 (2018).

51 Shang, H., Tong, H., Zhang, S., Chen, F. & Trinkaus, E. An early modern human from Tianyuan Cave, Zhoukoudian, China. Proc Natl Acad Sci U S A 104, 6573–6578 (2007).

52 Wang, F. G. et al. Innovative ochre processing and tool use in China 40,000 years ago. Nature 603, 284–289 (2022).

53 Yang, S. X. et al. Initial Upper Palaeolithic material culture by 45,000 years ago at Shiyu in northern China. Nat Ecol Evol 8, 552–563 (2024).

54 Li, F. et al. History, Chronology and Techno-Typology of the Upper Paleolithic Sequence in the Shuidonggou Area, Northern China. Journal of World Prehistory 32, 111–141 (2019).

55 Kim, K. J. et al. Radiocarbon Ages of Suyanggae Paleolithic Sites in Danyang, Korea. Radiocarbon 63, 1429–1444 (2021).

56 Fujita, M. et al. Advanced maritime adaptation in the western Pacific coastal region extends back to 35,000-30,000 years before present. Proceedings of the National Academy of Sciences of the United States of America 113, 11184–11189 (2016).

57 Sato, H. & Morisaki, K. On the beginning of the Japanese Upper Paleolithic_ A review of recent archaeological and anthropological evidence. Acta Anthroopologica Sinica (2022).

58 Ryan, W. B. F. et al. Global Multi-Resolution Topography synthesis. Geochemistry, Geophysics, Geosystems 10, Q03014 (2009).

59 Ho, C. K., Qi, G. Q. & Chang, C. H. A preliminary study of Late Pleistocene carnivore fossils from the Penghu Channel, Taiwan. Annual of Taiwan Museum 40, 1095–1224 (1997).

60 Asahara, M., Chang, C.-H., Kimura, J., Son, N. T. & Takai, M. Re-examination of the fossil raccoon dog (Nyctereutes procyonoides) from the Penghu channel, Taiwan, and an age estimation of the Penghu fauna. Anthropological Science 123, 177–184 (2015).

61 Tseng, Z. J. & Chang, C.-H. A study of new material of Crocuta crocuta ultima (Carnivora: Hyaenidae) from the Quaternary of Taiwan. Collection and Research 20, 9–19 (2007).

62 Sun, C.-H. et al. The southernmost Zalophus in the western North Pacific: the first pinniped (Carnivora, Pinnipedia) fossil from Taiwan. Journal of Vertebrate Paleontology 45 (2026).

63 Gao, J.-W. Penghu fauna. Journal of Marine Science (Geology Journal, Institute of Marine Science and Department of Marine Science, Chinese Culture University) 27, 123–132 (1982).

64 Ho, C. K., Qi, G. Q. & Chang, C. H. A preliminary study of late Pleistocene megafauna Cervus sp. from the Penghu Channel, Taiwan. Journal of the National Taiwan Museum 61, 1–16 (2008).

65 Chang, C. H. The first fossil record of a short-finned pilot whale (Globicephala macrorhynchus) from the Penghu Channel. Bulletin of National Museum of Natural Science 8, 73–80 (1996).

66 Tsai, C.-H., Fordyce, R. E., Chang, C.-H. & Lin, L.-K. Quaternary Fossil Gray Whales from Taiwan. Paleontological Research 18, 82–93 (2014).

67 Tsai, C. H. & Chang, C. H. A right whale (Mysticeti, Balaenidae) from the Pleistocene of Taiwan. Zoological Lett 5, 37 (2019).

68 Tan, K. On the fossil elephant remains in the Government Museum of Taiwan. Transactions of the Natural History Society of Formosa 25, 311–314 (1931).

69 Shikama, T., Otsuka, H. & Tomida, Y. Fossil Proboscidea from Taiwan. *Science Reports of the Yokohama National University*, Section II 22, 13–62 (1975).

70 Biswas, D. S., Chang, C.-H. & Tsai, C.-H. Land of the giants: Body mass estimates of Palaeoloxodon from the Pleistocene of Taiwan. Quaternary Science Reviews 336 (2024).

71. Shan, H.-y., Cheng, Y.-n. & Wu, X.-c. The first fossil skull of Alligator sinensis from the Pleistocene, Taiwan, with a paleogeographic implication of the species. Journal of Asian Earth Sciences 69, 17–25 (2013).

72 Liaw, Y. L. & Tsai, C. H. Taxonomic revision of Chinemys pani (Testudines: Geoemydidae) from the Pleistocene of Taiwan and its implications of conservation paleobiology. Anatomical record 306, 1501–1507 (2023).

73 Zhao, E. M. & Adler, K. Herpetology of China. Contributions to Herpetology No. 10., (Society for the Study of Amphibians and Reptiles, USA, 1993).

74 Liaw, Y.-L. & Tsai, C.-H. The first fossil of the extant leatherback sea turtle Dermochelys coriacea. Acta Palaeontologica Polonica 71, 267–271 (2026).

