## Supplementary Information for "Habitat and feeding ecology of a Denisovan from Late Pleistocene Taiwan"

This PDF file includes:
Supplementary Notes 1 to 6
and associated Supplementary Figures and Tables
Supplementary references

### Supplementary Note 1:
### Additional information and previous understanding of the Penghu fauna

The Taiwan Strait, with an average depth of approximately 60–70 m and numerous underwater gorges reaching depths of >100m, has undergone repeated emergence above sea level and submergence associated with Pleistocene sea-level fluctuations (**Extended Data Fig. 1**). Within this sea area, the Penghu Submarine Channel located between mainland Taiwan and the Penghu Islands has produced abundant vertebrate fossils, collectively known as the Penghu faunal assemblage (see **Extended Data Table 1** for major represented taxa). The situation is comparable to the English Channel in Europe or the Seto Inland Sea in Japan, where terrestrial vertebrates inhabited the exposed land during the Pleistocene glaciations.

The discovery of vertebrate fossils from the Penghu seafloor traces back to almost a century ago, when a well-preserved mandible of the large elephantid *Palaeoloxodon* was described based on a museum collection[1]. However, these submarine fossils received little attention for the next several decades. Detailed study of the Penghu faunal assemblage began around the end of the 20th century[2-5] and research was further activated in the past two decades[6-8], supported by the increasing recovery of fossils through commercial trawler fishing during the latter half of the 20th century.

Additional information on the Penghu assemblage was gathered by one of us (C.-H. C.), through repeated interviews with fishermen in southern Taiwan, including one who had 34 years (1986–2020) of experience in commercial fishing and incidental recovery of vertebrate fossils from the seabed. According to this information, such fossils are dredged from the area ranging from 119°47′E to 119°54′E and 23°00′N to 23°50′N (**Extended Data Fig. 1**). The fishing vessels employ bottom trawl nets approximately 4–6 m in width with a mesh size of approximately 9 × 9 cm. Trawling operations are typically conducted at water depths of 80 to 160 m. After deployment, the trawl net is towed with the prevailing current at approximately 4–5 knots (nautical mile/h) for about 1 hour. Consequently, each trawling operation covers approximately 3–5 nautical miles (ca. 5.5–9.3 km) across the seafloor. Given the relatively large mesh size of the trawl nets, fossil recovery is strongly biased toward larger specimens, whereas small vertebrate remains are unlikely to be retained. In addition, because the trawl is towed continuously over several kilometers, the precise provenance and stratigraphic horizon of individual fossils cannot be determined, and each recovered specimen may have originated from any location along the trawled path.

The majority of the Penghu assemblage consists of extinct species or species no longer present in the modern Taiwanese ecosystem. Previous analyses of carbon and oxygen isotopes suggested that *Palaeoloxodon* from this area inhabited a C4-dominated ecosystem with relatively abundant water resources, similar to the modern African savannah[9] (see also an illustrated landscape reconstruction in this reference). The open, savannah-like environmental model is consistent with the occurrence of horse (*Equus*) and hyena (*Crocuta*) in Penghu fauna, although some taxa (*Nyctereutes procyonoides*, *Ursus arctos, Panthera tigris*, etc.) suggest the presence of woodlands. In combination with the results of the oxygen isotope analysis[9], the topography of the area strongly suggests the presence of a monsoon-fed river system, which is consistent with the existence of some freshwater taxa such as alligator (*Alligator*) and pond turtle (*Mauremys*), as well as milu (*Elaphurus*) and water buffalo (*Bubalus*). In summary, the so far available fragmentary evidence at least suggests the presence of open and wet environments as well as patchy or temporary woodlands in the Penghu region.

Dating the Penghu fossils has been challenging primarily because their stratigraphic contexts are unknown. The previous attempts of direct dating by the uranium-series method were complicated by interaction with uranium from the sea water[10,11]. Therefore, many researchers have relied on biochronological comparisons with fossil sequences reported from the continental Asia[10,12], which generally suggests Middle to Late Pleistocene ages. For example, the occurrence of *Crocuta* suggests that the Penghu fauna is no older than the late Middle Pleistocene[10]. However, given the uncertainty of geological horizons for each fossil specimen, some researchers prefer to assign a broader Middle to Late Pleistocene age (approximately 0.78-0.01 Ma)[8,9,13].

**Supplementary Note 2:**
**Research history of Penghu 2 and 3**

Soon after we (C.-H.C., Y.K., Masanaru Takai and Reiko T. Kono) started our work on the Penghu 1 mandible that derived from an antique shop in Tainan City, we identified Penghu 2 and 3 in October 2010 at the National Museum of Natural Science of Taiwan located in Taichung City (hereafter 'NMNS Taiwan'). These hominin leg bones were among a large Penghu fossil collection collected by Mr. Li-Ren Hou, a local artist, who donated it to the NMNS Taiwan, in 2011. Following this discovery, we initiated a stepwise research program in 2012 to analyze these hominin leg bones as summarized below, alongside our investigations of Penghu 1, which were published in 2015[10] and 2025[14].

In 2012, we cut off a small piece of bone from the distal end of each leg bone (as well as the central part of the Penghu 1 mandible) using a thin diamond cutter at the National Museum of Nature and Science, Tokyo (hereafter 'NMNS Tokyo'). We submitted the separated fragments to laser- ablation U-series dating, the results of which were recently published in ref. 11. Based on the bone powder generated from this cutting, we also examined organic contents and obtained negative and positive results for Penghu 2 and 3, respectively. In 2015, at NMNS Taiwan, we took additional bone samples of Penghu 2 and 3 by drilling the sections created by the 2012 work. We also collected bone samples from non-human mammalian fossils from the Penghu Channel at that opportunity. Based on these samples, in 2017, we obtained the radiocarbon dates we reported in the present paper. This positive result for Penghu 3 prompted us to collect additional bone samples from these fossils in Tokyo for palaeoproteomic analysis and ancient DNA extraction in 2019 and 2024. The palaeoproteomic results are reported in ref. 15, and the outcome of the ancient DNA analyses will be reported elsewhere by a team of The University of Tokyo and NMNS Taiwan.

For morphological analyses, we performed pct. scans of the two leg bones at The University Museum, The University of Tokyo, in 2024. Because we found that it is difficult to digitally separate the attached nodules from the bone surfaces due to ambiguous boundaries in the CT scan, we physically removed these nodules at the NMNS Taiwan in 2025[15].

**Supplementary Note 3:**
**Prescreening**

A total of 72 skeletal specimens, consisting of three *Homo* specimens and 69 faunal specimens, were selected for the prescreening (**Supplementary Table 3_1**). According to a previous study, specimens with a bone nitrogen concentration (%N) of 0.76 wt% or higher and an atomic carbon-to-nitrogen ratio (C/N) of 17 or lower are considered likely to retain endogenous collagen[16]. Among the Penghu specimens, only two—F051713 (Penghu 3) and F027088 (*Bubalus*)— satisfied the former criterion (bone %N = 1.7 wt% and 0.7 wt%, respectively; **Supplementary Table 3_1**). In contrast, 16 specimens, including F051713 and F027088, satisfied the latter criterion (bone C/N ratio ≤ 17). Because collagen can sometimes be extracted even from bones or dentine that do not meet these criteria, the prescreening threshold was relaxed in this study, and 32 specimens with bone %N of ≥0.2 wt% were selected for collagen extraction. Among the 72 specimens subjected to prescreening, Penghu 3 had the highest bone %N (1.7 wt%) and the lowest bone C/N ratio (5.0), and was therefore inferred preserve collagen best. The bone %N values for Penghu 1 and Penghu 2 fell below the detection limit, indicating they retained almost no extractable collagen and were not used for collagen extraction.

Collagen extraction was performed on 32 bone and dentine specimens that met the prescreening criterion of bone %N ≥ 0.2% (**Supplementary Table 3_2**). For eight specimens with larger sample amounts, up to three independent collagen extractions were performed. Overall preservation was poor, and no gelatin was recovered in 9 of the 32 specimens (9 of 40 extractants). The median gelatin yield was 0.2% among the 31 extractants with recovered gelatin.

Based on gelatin yield and elemental concentrations, we selected four specimens (F051713 [Penghu 3], F027088, F027093, and F051471) for further analyses (**Supplementary Table 3_2**). Four specimens (F051713, F027088, F051471, and F051688) met the empirical criterion of gelatin yield (wt%) ≥ 1%, which is commonly used to select well-preserved samples for radiocarbon dating[17]. Five specimens (F051713, F027088, F027093, F051471, and P201508-6) satisfied the criterion of the elemental C/N ratio of gelatin (2.9–3.4)[18], used to confirm collagen integrity. Three specimens that met both criteria (F051713, F027088, and F051471) also met the criteria of carbon and nitrogen elemental concentrations of gelatin (%C > 13%, %N > 4.8%)[17]. Specimen F051688 had yields of 1.3% and 0.3%, thus passing the yield criterion, but its C/N ratios of 4.6 and 3.8 failed the C/N criterion. Therefore, F051688 was excluded from most discussions, although one of the extractants (PCO-4190) of this specimen was included and treated separately in the CSIA analyses (see Supplementary Note S5). Specimen F027093 (PCO-1071) did not meet the yield criterion (0.8%) but did satisfy the C/N ratio criterion (3.1), and its gelatin %C and %N values (36.6% and 13.8%, respectively) fell within the acceptable ranges. Therefore, F027093 was included in further analyses. Although P201508-6 satisfied the criterion of C/N ratio (3.4), its yield was very low (0.1%), and its gelatin %C and %N values (5.2% and 1.8%, respectively) were below the acceptable range.

Therefore, P201508-6 was considered to have insufficient collagen integrity and was excluded from subsequent analyses.

Based on the results of collagen extraction and elemental concentrations, the prescreening metrics for the Penghu fossils were retrospectively evaluated. In specimens with the successful collagen extraction and acceptable collagen quality, as indicated by elemental concentrations and C/N ratios[17,18] ($n = 4$), bone %N was ≥0.4 wt%, and the bone C/N ratio was ≤7.9 (**Supplementary Table 3_1**). Because these prescreening thresholds are also influenced by environmental factors such as the degree of post-mortem mineralization of skeletal materials, they need to be calibrated for individual sites.

**Supplementary Table 3_1**. **Prescreening results of the subject Penghu fossils.** The identifiers shown in the "Number" column are identical to those in the Supplementary Tables 3_2. and 4_1.

| **No.** | **NMNS ID** | **Taxon** | **Skeletal element** | **Bone C concentration (wt%)** | **Bone N concentration (wt%)** | **Bone atomic C/N ratio** |
|---|---|---|---|---|---|---|
| 1 | F051911 (Penghu 1) | *Homo* | Mandible | 2.6 | N.D. | N.D. |
| 2 | F051716 (Penghu 2) | *Homo* | Femur | 2.7 | N.D. | N.D. |
| 3 | F051713 (Penghu 3) | *Homo* | Tibia | 7.3 | 1.7 | 5.0 |
| 4 | F026918 | *Palaeoloxodon* | Humerus | 1.8 | N.D. | N.D. |
| 5 | F026951 | *Palaeoloxodon* | Humerus | 1.8 | N.D. | N.D. |
| 6 | F027004 | *Palaeoloxodon* | Rib | 2.7 | N.D. | N.D. |
| 7 | F027005 | *Palaeoloxodon* | Rib | 1.9 | N.D. | N.D. |
| 8 | F027006 | *Palaeoloxodon* | Rib | 2.1 | 0.1 | 24.5 |
| 9 | F027026 | *Bubalus* | Humerus | 2.6 | 0.1 | 30.3 |
| 10 | F027033 | *Bubalus* | Tibia | 2.1 | 0.1 | 24.5 |
| 11 | F027034 | *Bubalus* | Tibia | 2.2 | 0.3 | 8.6 |
| 12 | | | | - | - | - |
| 13 | | | | - | - | - |
| 14 | F027088 | *Bubalus* | Tibia | 3.1 | 0.7 | 5.2 |
| 15 | | | | - | - | - |
| 16 | F027091 | *Bubalus* | Tibia | 2.0 | 0.1 | 23.3 |
| 17 | | | | 1.9 | N.D. | N.D. |
| 18 | F027093 | *Bubalus* | Humerus | 2.7 | 0.4 | 7.9 |
| 19 | | | | - | - | - |
| 20 | | | | - | - | - |
| 21 | F028047 | *Palaeoloxodon* | Tibia | 1.5 | N.D. | N.D. |
| 22 | F028061 | *Palaeoloxodon* | Limb | 2.1 | N.D. | N.D. |
| 23 | F028941 | *Palaeoloxodon* | Femur | 2.3 | 0.1 | 26.8 |
| 24 | F051470 | *Cervus* | Mandible | 1.6 | 0.1 | 18.7 |
| 25 | | | | - | - | - |
| 26 | F051471 | *Cervus* | Mandible | 3.0 | 0.5 | 7.0 |
| 27 | | | | - | - | - |
| 28 | | | | - | - | - |
| 29 | F051687 | *Equus* | Tibia | 1.7 | 0.1 | 19.8 |
| 30 | F051688 | *Equus* | Tibia or femur | 3.2 | 0.5 | 7.5 |

| | | | | | | |
|---|---|---|---|---|---|---|
| 31 | | | | - | - | - |
| 32 | F051689 | *Equus* | Tibia | 2.4 | 0.1 | 28.0 |
| 33 | F051690 | *Equus* | Tibia | 1.5 | 0.1 | 17.5 |
| 34 | F051693 | *Equus* | Radius and ulna | 1.6 | N.D. | N.D. |
| 35 | F051694 | *Equus* | Radius | 2.1 | N.D. | N.D. |
| 36 | F051695 | *Equus* | Tibia | 2.1 | N.D. | N.D. |
| 37 | F051702 | *Equus* | Tibia | 2.6 | N.D. | N.D. |
| 38 | F051705 | *Equus* | Maxilla | 2.4 | 0.3 | 9.3 |
| 39 | | | | - | - | |
| 40 | F051706 | *Equus* | Mandible | 2.4 | N.D. | N.D. |
| 41 | F051717 | *Sus* | Cranium | 2.5 | N.D. | N.D. |
| 42 | F051718 | *Sus* | Humerus | 1.6 | 0.1 | 18.7 |
| 43 | F051719 | *Sus* | Humerus | 1.2 | N.D. | N.D. |
| 44 | F051720 | *Sus* | Humerus | 2.2 | 0.1 | 25.7 |
| 45 | F051721 | *Sus* | Humerus | 1.4 | 0.1 | 16.3 |
| 46 | F27072 | *Bubalus* | Tibia | 1.8 | 0.2 | 10.5 |
| 47 | F27073 | *Bubalus* | Humerus | 1.6 | N.D. | N.D. |
| 48 | F27074 | *Bubalus* | Humerus | 2.4 | 0.2 | 14.0 |
| 49 | P201508-1 | Euungulata | Vertebrate | 2.1 | 0.1 | 24.5 |
| 50 | P201508-2 | *Equus* | Metatarsus | 1.8 | N.D. | N.D. |
| 51 | P201508-3 | Euungulata | Phalanx | 1.7 | N.D. | N.D. |
| 52 | P201508-4 | Euungulata | Scapula | 1.9 | N.D. | N.D. |
| 53 | P201508-5 | *Bubalus* | Molar (left $M_3$) | 2.0 | 0.1 | 23.3 |
| 54 | P201508-6 | *Bubalus* | Molar | 3.9 | 0.3 | 15.2 |
| 55 | P201508-7 | *Bubalus* | Upper molar | 1.2 | 0.1 | 14.0 |
| 56 | P201508-8 | Proboscidea | Bone fragment | 1.9 | 0.1 | 22.2 |
| 57 | P201508-9 | Proboscidea | Bone fragment | 2.5 | 0.1 | 29.2 |
| 58 | P201508-10 | N.D. | Bone fragment | 2.3 | 0.1 | 26.8 |
| 59 | P201508-11 | Proboscidea | Bone fragment | 1.6 | N.D. | N.D. |
| 60 | P201508-12 | *Palaeoloxodon?* | Bone fragment | 3.0 | 0.3 | 11.7 |
| 61 | P201508-13 | N.D. | Bone fragment | 2.0 | N.D. | N.D. |
| 62 | P201508-14 | N.D. | Bone fragment | 2.1 | N.D. | N.D. |
| 63 | P201508-15 | *Bubalus* | Bone fragment | 1.9 | 0.1 | 22.2 |
| 64 | P201508-16 | *Bubalus* | Bone fragment | 2.3 | 0.1 | 26.8 |
| 65 | P201508-17 | *Bubalus* | Bone fragment | 2.2 | 0.2 | 12.8 |
| 66 | P201508-19A | N.D. | Bone fragment | 1.9 | 0.1 | 22.2 |

| | | | | | | |
|---|---|---|---|---|---|---|
| 67 | P201508-19B | N.D. | Bone fragment | 2.3 | N.D. | N.D. |
| 68 | P201508-20 | *Bubalus* | Molar (left $M_3$) | 1.7 | 0.1 | 19.8 |
| 69 | P201508-21 | Proboscidea | Molar fragment | 0.7 | 0.1 | 8.2 |
| 70 | P201508-22 | N.D. | Bone fragment | 2.1 | 0.1 | 24.5 |
| 71 | P201508-23 | Cervidae | Antler | 2.3 | 0.3 | 8.9 |
| 72 | P201508-24 | *Bubalus* | Humerus | 2.2 | N.D. | N.D. |
| 73 | P201508-25 | N.D. | Bone fragment | 2.3 | 0.1 | 26.8 |
| 74 | P201508-26 | N.D. | Bone fragment | 1.7 | N.D. | N.D. |
| 75 | P201508-27 | *Bubalus* | Metacarpal | 2.3 | 0.1 | 26.8 |
| 76 | P201508-28 | N.D. | Bone fragment | 2.6 | 0.1 | 30.3 |
| 77 | P201508-29 | N.D. | Bone fragment | 2.2 | 0.1 | 25.7 |
| 78 | P201508-30 | N.D. | Bone fragment | 2.4 | N.D. | N.D. |
| 79 | N.D. | *Bubalus* | Limb shaft | 1.5 | N.D. | N.D. |
| 80 | N.D. | Euungulata | Limb shaft | 1.3 | N.D. | N.D. |
| 81 | N.D. | *Bubalus* | Limb shaft | 2.3 | N.D. | N.D. |
| 82 | N.D. | Euungulata | Limb shaft | 1.6 | N.D. | N.D. |
| 83 | N.D. | Euungulata | Limb shaft | 2.8 | N.D. | N.D. |

**Supplementary Table 3_2**. **Collagen extraction and bulk stable isotopic results of the subject Penghu fossils.** The identifiers shown in the "Number" column are identical to those in the Supplementary Tables 3_1 and 4_1.

| **Number** | **NMNS ID** | **Preparation No.** | **Yield (wt%)** | **Collagen C concentration (wt%)** | **Collagen N concentration (wt%)** | **Collagen atomic C/N ratio** | **$\delta^{13}C$ (‰)** | **$\delta^{15}N$ (‰)** |
|---|---|---|---|---|---|---|---|---|
| 3 | F051713 | PCO-1163 | 6.7 | 43.3 | 15.6 | 3.2 | -6.7 | 11.4 |
| 4 | F026918 | PCO-1082 | 0.1 | - | - | - | - | - |
| 5 | F026951 | PCO-1078 | 0.0 | - | - | - | - | - |
| 6 | F027004 | PCO-1080 | 0.1 | 11.2 | 1.7 | 7.5 | -18.8 | 3.3 |
| 7 | F027005 | PCO-1079 | 0.0 | - | - | - | - | - |
| 8 | F027006 | PCO-1081 | 0.1 | 14.6 | 2.2 | 7.6 | -23.0 | 2.2 |
| 9 | F027026 | PCO-1075 | 0.1 | - | - | - | - | - |
| 10 | F027033 | PCO-1076 | 0.0 | - | - | - | - | - |
| 11 | F027034 | PCO-1077 | 0.3 | 8.7 | 2.1 | 4.8 | -9.9 | 0.7 |
| 12 | | PCO-1161 | 0.2 | 18.1 | 5.6 | 3.8 | -8.9 | 3.0 |

| | | | | | | | | |
|---|---|---|---|---|---|---|---|---|
| 13 | | PCO-1742 | 0.1 | - | - | - | - | - |
| 14 | F027088 | PCO-1074 | 1.7 | 38.6 | 14.7 | 3.1 | -5.5 | 5.8 |
| 15 | | PCO-1741 | 0.0 | - | - | - | - | - |
| 16 | F027091 | PCO-1073 | 0.1 | 24.0 | 4.9 | 5.8 | -16.5 | 3.3 |
| 17 | | PCO-1072 | 0.1 | - | - | - | - | - |
| 18 | F027093 | PCO-1071 | 0.8 | 36.6 | 13.8 | 3.1 | -7.7 | 9.8 |
| 19 | | PCO-1740 | 0.3 | 27.7 | 9.7 | 3.3 | -6.9 | 7.0 |
| 20 | | PCO-4188 | 0.2 | 22.3 | 7.1 | 3.6 | -8.1 | 8.8 |
| 21 | F028047 | PCO-1084 | 0.0 | - | - | - | - | - |
| 22 | F028061 | PCO-1083 | 0.0 | - | - | - | - | - |
| 23 | F028941 | PCO-1085 | 0.2 | 4.7 | 0.6 | 9.2 | -20.9 | 5.1 |
| 24 | F051470 | PCO-1066 | 0.1 | - | - | - | - | - |
| 25 | | PCO-1743 | 0.0 | - | - | - | - | - |
| 26 | F051471 | PCO-1067 | 2.4 | 40.9 | 15.6 | 3.1 | -10.1 | 8.4 |
| 27 | | PCO-1739 | 2.2 | 41.4 | 14.5 | 3.3 | -11.4 | 7.4 |
| 28 | | PCO-4189 | 1.0 | 34.3 | 11.3 | 3.5 | -10.7 | 6.9 |
| 30 | F051688 | PCO-1735 | 0.3 | 35.5 | 11.0 | 3.8 | -8.3 | 7.8 |
| 31 | | PCO-4190 | 1.3 | 16.2 | 4.1 | 4.6 | -7.6 | 6.3 |
| 33 | F051690 | PCO-1736 | 0.0 | - | - | - | - | - |
| 38 | F051705 | PCO-1737 | 0.3 | 24.9 | 6.5 | 4.5 | -10.8 | 10.8 |
| 39 | | PCO-4191 | 0.6 | 10.0 | 0.1 | 170.7 | -13.6 | 7.2 |
| 45 | F051721 | PCO-1738 | 0.0 | - | - | - | - | - |
| 46 | F27072 | PCO-1068 | 0.3 | - | - | - | - | - |
| 47 | F27073 | PCO-1069 | 0.1 | 29.1 | 5.9 | 5.7 | -19.3 | 4.7 |
| 48 | F27074 | PCO-1070 | 0.2 | 31.3 | 8.9 | 4.1 | -11.9 | 6.3 |
| 49 | P201508-1 | PCO-1167 | 0.2 | 37.1 | 9.4 | 4.6 | -11.7 | 6.4 |
| 54 | P201508-6 | PCO-1168 | 0.1 | 5.2 | 1.8 | 3.4 | -24.3 | -0.8 |
| 60 | P201508-12 | PCO-1169 | 0.6 | 33.9 | 10.3 | 3.8 | -13.4 | 7.1 |
| 65 | P201508-17 | PCO-1170 | 0.1 | 12.8 | 4.1 | 3.7 | -9.4 | 3.9 |
| 71 | P201508-23 | PCO-1172 | 0.2 | 26.5 | 6.6 | 4.7 | -17.0 | 3.9 |

### Supplementary Note 4:
### Radiocarbon measurement

#### Improved collagen quality by ultrafiltration

Because ultrafiltration (UF) concentrates high–molecular-weight collagen and removes exogenous contaminants, it yields more accurate ages, particularly for samples older than 40 ka[19]. The variability of $^{14}C$ ages obtained without UF, and differences between $^{14}C$ ages with and without UF were evaluated to assess the effect of UF on the Penghu fossils.

To examine the variability in $^{14}C$ ages without UF, AMS measurements were performed on collagen samples prepared without UF from three independent extractions of two faunal specimens (F027093 and F051471) (**Supplementary Table 4_1**; **Extended Data Table 2**). For F027093, one replicate had a low collagen yield, and the result of an ultra-micro (170 µgC) AMS measurement fell below the quantification limit (>30,000 BP); this replicate was excluded from further analyses. The remaining two replicates yielded conventional radiocarbon ages (CRA) of 34,804 ± 135 BP (TKA-16642) and 43,149 ± 282 BP (TKA-22245), differing by 8345 $^{14}C$ years (**Supplementary Table 4_1**). For F051471, three replicates yielded CRA of 36,735 ± 164 $^{14}C$ years BP (TKA-16641), 40,650 ± 261 $^{14}C$ years BP (TKA-29399), and 39,708 ± 209 $^{14}C$ years BP (TKA-22246), spanning 3915 $^{14}C$ years (**Supplementary Table 4_1**). These large discrepancies suggest contamination by exogenous carbon compounds in varying proportions thereby reducing the precision of the $^{14}C$ age estimates in samples without UF.

The effectiveness of UF in removing such exogenous carbon compounds was evaluated by comparing CRA with and without UF across three extractants. For extractant PCO-4188 from specimen F027093 and extractants PCO-1739 and PCO-4189 from specimen F051471, UF yielded CRAs that were 1679 $^{14}C$ years younger, 812 $^{14}C$ years older, and 446 $^{14}C$ years older, respectively (**Supplementary Table 4_1**; **Extended Data Table 2**). The difference between the CRAs of two extractants from the same F051471 specimen was 1308 $^{14}C$ years for UF-treated samples, smaller than that for non-UF-treated samples (3915 $^{14}C$ years). The fact that UF produced younger, rather than systematically older, $^{14}C$ ages for some extractants indicates that the shifts cannot be explained by an artefactual contribution of dead carbon retained on the UF membrane. Instead, the convergence of CRAs among extractants from different specimens after UF suggests that UF successfully removed exogenous carbon compounds and yielded more accurate CRAs derived from endogenous collagen.

The $^{14}C$ ages of Penghu 3 were compared with and without UF. The UF-treated collagen yielded a CRA of 42,399 ± 313 $^{14}C$ years BP (TKA-30498), which is 1756 $^{14}C$ years older than the CRA of non-UF-treated gelatin of 40,643 ± 204 $^{14}C$ years BP (TKA-16645) (**Supplementary Table 4_1**). This corresponds to a difference of 1325 ± 403 years in calibrated age (**Extended Data Fig. 2**). The UF-treated collagen from Penghu 3 yielded CRAs that closely match those obtained from UF-treated faunal bone collagen (differing by 929–2245 $^{14}C$ years in CRA). In addition, Penghu 3 consistently showed outstanding quality in both prescreening metrics and collagen-based indicators (**Supplementary Tables 3_1 and 3_2**). Taken together, these observations support the reliability of the $^{14}C$ age estimate of Penghu 3.

**Evaluation of the $^{14}C$ detection limit**

Because the $^{14}C$ age for Penghu 3 without UF lay very close to the quantification limit of usual measurements at UMUT (**Supplementary Table 4_1**), the AMS measurement conditions were adapted for old samples with UF. Specifically, for the three UF-treated samples, the number of concurrently measured standards was increased (five HOxII-2 standards and three Wako-OX blanks) to verify that the measured values ($^{14}C$/ $^{12}C$) lay above the quantification limit (**Extended Data Fig. 3**). Wako-OX is a $^{14}C$-free material whose measured values reflect the minimum background level resulting from unavoidable $^{14}C$ contamination during $CO_2$ production, graphitization, and AMS measurement. The $^{14}C$ concentrations of multiple Wako-OX measurements were stable, indicating that the measurement background is well constrained. The $^{14}C$ concentrations obtained for the UF-treated samples consistently exceed the quantification limit (average + 10 standard deviations with the Wako-OX measurements). When the measurement results for IAEA-C6 and IAEA-C8 are calibrated against those for HOxII and Wako-OX, they correspond very closely to their respective consensus values (**Supplementary Table 4_2**), implying that, although the Penghu fossils lie close to the quantification limit, the precise $^{14}C$ concentrations were reliably measured in our AMS system.

**Supplementary Table 4_1. Radiocarbon results of the subject Penghu fossils.** The identifiers shown in the "Number" column are identical to those in the Supplementary Tables 3_1 and 3_2. Results of the same extractants before and after applying ultrafiltration (UF) are shown for some samples. Data of samples with measured $^{14}C$ contents below the quantification limit are shown in italics and with an asterisk in the column "Amount of graphite".

| Number | NMNS ID | Prep. No. | Before UF | | | After UF | | |
|---|---|---|---|---|---|---|---|---|
| | | | AMS lab ID | CRA (years BP) | Amount of graphite (mg) | AMS lab ID | CRA (years BP) | Amount of graphite (mg) |
| 3 | F051713 | PCO-1163 | TKA-16645 | 40,643 ± 204 | 0.97 | TKA-30498 | 42,399 ± 313 | 0.94 |
| 14 | F027088 | PCO-1074 | TKA-16643 | 35,002 ± 143 | 1.1 | - | - | - |
| 18 | F027093 | PCO-1071 | TKA-16642 | 34,804 ± 135 | 1.02 | - | - | - |
| 19 | | *PCO-1740* | *TKA-29469* | *>30,000* | *0.17** | - | - | - |
| 20 | | PCO-4188 | TKA-22245 | 43,149 ± 282 | 1.19 | TKA-30499 | 41,470 ± 297 | 0.76 |
| 26 | F051471 | PCO-1067 | TKA-16641 | 36,735 ± 164 | 1.16 | - | - | - |
| 27 | | PCO-1739 | TKA-29399 | 40,650 ± 261 | 1.01 | TKA-31973 | 41,462 ± 343 | - |
| 28 | | PCO-4189 | TKA-22246 | 39,708 ± 209 | 0.93 | TKA-30500 | 40,154 ± 252 | 0.79 |
| 30 | F051688 | *PCO-1735* | *TKA-17737* | *> 39,000* | *0.11** | - | - | - |
| 31 | | PCO-4190 | TKA-22247 | 28,049 ± 92 | 0.89 | - | - | - |
| 38 | F051705 | *PCO-1737* | *TKA-17738* | *23,802 ± 176* | *ca.0.15** | - | - | - |
| 60 | P201508-12 | PCO-1169 | TKA-16644 | 36,737 ± 223 | 0.63 | - | - | - |

**Supplementary Table 4_2. Summary of AMS $^{14}$C results for standards, blanks, and UF-treated samples.** Values with an asterisk were used for calibration. References for the Source column correspond to refs. [20,21].

| **Sample** | **No. of targets** | **No. of runs** | **$^{14}$C/$^{12}$C** | | **Measured F$^{14}$C * 100** | | **Consensus F$^{14}$C * 100** | | **Source** |
|---|---|---|---|---|---|---|---|---|---|
| | | | Average | SD | Average | SD | Average | SD | |
| NIST SR 4990C (HOxII) | 5 | 35 | 1.53909E-12 | 9.90958E-15 | 134.066* | 0.109 | 134.066 | 0.043 | Ref. 20 |
| Process blank (Wako-OX) | 3 | 21 | 1.94651E-15 | 2.95565E-16 | 0.173* | 0.005 | 0 | - | |
| Machine blank (Alpha1) | 1 | 7 | 6.70E-16 | 1.78091E-16 | | | | | |
| IAEA-C6 | 1 | 7 | 1.75176E-12 | 1.24607E-14 | 150.61 | 0.37 | 150.61 | 0.11 | Ref. 21 |
| IAEA-C8 | 1 | 7 | 1.73281E-13 | 2.07286E-15 | 14.96 | 0.08 | 15.03 | 0.17 | Ref. 21 |
| TKA-30498 | 1 | 7 | 7.90E-15 | 4.27228E-16 | | | | | |
| TKA-30499 | 1 | 7 | 8.60882E-15 | 5.36953E-16 | | | | | |
| TKA-30500 | 1 | 7 | 9.78673E-15 | 6.71062E-16 | | | | | |

### Supplementary Note 5:
### Bulk stable isotope analysis

#### Reproducibility of the bulk stable isotope ratios

To assess the precision of bulk collagen stable isotope ratios calibrated against three working standards (**Supplementary Table 5**), the reproducibility of repeated collagen extractions and isotope analyses was investigated. Up to three independent replicate collagen extractions and isotope ratio mass spectrometry (IRMS) measurements were conducted for five specimens (F027034, F027093, F051471, F051688, F051705). Most specimens did not yield sufficient collagen, and two of them, F027093 (*Bubalus*) and F051471 (*Cervus*), with acceptable atomic C/N ratios in the extracted collagen samples (**Supplementary Table 3_2**), were included in the analysis. Across three replicate extractions and measurements for specimens F027093 and F051471, the mean $\delta^{13}C$ values were −7.6 ± 0.6‰ and −10.7 ± 0.7‰, and the mean $\delta^{15}N$ values were 8.5 ± 1.4‰ and 7.6 ± 0.8‰, respectively, with one standard deviation (**Supplementary Table 3_2**). These dispersions exceeded the instrumental uncertainty of < ±0.1‰ determined for the IRMS, indicating that some influence of exogenous contaminants on collagen samples cannot be entirely ruled out. However, $\delta^{13}C$ values of C3 and C4 plants typically differ by about 10‰[22], which is a much larger difference than the observed dispersion. Thus, the $\delta^{13}C$ value of −6.7‰ obtained for Penghu 3 is clearly consistent with a C4-dominated ecosystem (**Fig. 3**). In addition, Penghu 3 shows a $\delta^{15}N$ value 3.3‰ higher than that of F027093 (*Bubalus*), which has the highest nitrogen isotope ratio among the Penghu herbivores (**Table 2**). Compound-specific isotope analysis (CSIA), which largely removes the effects of exogenous contaminants, also independently indicates a high trophic position of 2.9 for Penghu 3 (**Supplementary Note 6**). Taken together, these observations show that the observed dispersion in bulk collagen stable isotope measurements has a negligible effect on our conclusions regarding the dietary niche of Penghu 3.

#### Aquatic resource use not supported by bulk isotope analysis

In general, relatively high bulk collagen $\delta^{13}C$ and $\delta^{15}N$ values are sometimes interpreted as evidence of substantial consumption of aquatic resources, including marine and freshwater resources[23,24]. Penghu 3 likewise exhibits comparatively high bulk $\delta^{13}C$ and $\delta^{15}N$ values (**Fig. 3**); however, as outlined below, an explanation based on intensive exploitation of aquatic resources appears unlikely and is better explained by strong carnivory within a terrestrial C4-dominated ecosystem.

First, the $\delta^{13}C$ value of Penghu 3 (−6.8‰) falls within the range (−5.5‰ to −10‰) observed for two *Bubalus* and one *Cervus* specimens from Penghu (**Fig. 3**) that yielded closely similar radiocarbon ages (**Fig. 2**). The $\delta^{13}C$ overlap suggests that all of these taxa originated from the same ecosystem, with C4 plants as primary producers (**Fig. 3**). This inference is consistent with their $\delta^{15}N_{Phe}$ values obtained in CSIA (**Supplementary Note 6**).

Moreover, along with the other terrestrial herbivores from Penghu, the $\delta^{13}C$ value of Penghu 3 is too high to be explained by aquatic inputs alone. Although collagen isotopic data for marine fauna from the Taiwan Strait are unavailable, Penghu 3 has $\delta^{13}C$ values more than

5‰ higher than those typically observed for marine mammals (−13.3‰ to −12.2‰), offshore pelagic fish (−14.7‰ to −12.7‰), salmonids (−16.3‰ to −14.5‰), freshwater fish (−19.0‰ to −16.6‰), and shellfish (−11.8‰ to −8.7‰) from nearby Japanese archipelago[23].

In modern forest ecosystems, a trophic enrichment of approximately 3–5‰ in bulk $\delta^{15}N$ values is commonly observed between herbivores and carnivores[25]. Using the mean $\delta^{15}N$ values of the three Penghu herbivores (**Table 2**), the expected $\delta^{15}N$ range for carnivores in this ecosystem is 10.2–12.2‰, which agrees well with the $\delta^{15}N$ value of Penghu 3 (11.4‰). Thus, the elevated bulk collagen $\delta^{15}N$ value of Penghu 3 can be explained by a high trophic level and strong faunivory, without assuming a contribution from aquatic resources.

Sulfur isotope ratios in collagen could, in principle, help detect dietary inputs from aquatic resources[24]. However, such analyses require 5–10 mg of collagen, which is more than an order of magnitude greater than the amount needed for bulk carbon and nitrogen stable isotope analysis. Given that the existing data provide no compelling evidence of significant aquatic resource use and that the preservation of precious Denisovan skeletal materials must take priority, we did not undertake sulfur isotope analysis in this study.

**Supplementary Table 5. Details of the working standards used for bulk stable isotope analysis.**

| **Compound** | **Lot number** | **$\delta^{13}C$ (‰)** | **$\delta^{15}N$ (‰)** | **Caliblants of $\delta^{13}N$** | **Caliblants of $\delta^{15}N$** |
|---|---|---|---|---|---|
| L-Alanine | AZ100 SS09 | -19.6 | 8.72 | NBS19, IAEA-C6 | IAEA N1, IAEA N2 |
| L-Histidine | AZ1Z0M6M9675 | -11.4 | -7.58 | NBS19, IAEA-C6 | IAEA N1, IAEA N2 |
| Glycine | AZ300M9R2283 | -32.3 | 1.12 | NBS19, IAEA-C6 | IAEA N1, IAEA N2 |

### Supplementary Note 6:
### Compound-specific stable nitrogen isotope analysis of amino acids

#### Validity of the selected source and trophic amino acid pairs

In compound-specific isotope analysis (CSIA) for dietary reconstruction, the trophic position (TP) of an animal can be calculated by comparing the $\delta^{15}N$ values of individual amino acids that undergo strong trophic enrichment (trophic amino acids) with those that show minimal ecological enrichment and retain the producers' baseline (source amino acids). This trait of CSIA enables calculation of an animal's more accurate TP while avoiding the confounding effects of variation in the isotopic baseline, unlike bulk stable isotope analysis[26,27].

Calculation of TP is based on the observation that the $\delta^{15}N$ values of trophic amino acids systematically increase at each step of the food chain, relative to the baseline difference between trophic and source amino acids in primary producers[28]. Several trophic–source amino acid pairs have been proposed for CSIA-based TP estimates, including glutamic acid (Glu)–phenylalanine (Phe) and proline (Pro)–lysine (Lys)[29]. Among these, the approach that calculates TP from the difference in nitrogen isotope ratios between Glu and Phe ($\Delta^{15}N_{Glu–Phe}$) is so far the most widely used in ecological and archaeological CSIA studies[30-35]. Accordingly, we adopted the Glu–Phe pair as the trophic–source pair for TP calculation and estimated TP from their nitrogen isotope ratios.

Other amino acid pairs proposed in previous studies cannot be measured with sufficiently high precision in the analytical system used in this study. Lys has been suggested to yield more consistent TP estimates than Phe, because its $\delta^{15}N$ values, as a source amino acid, show smaller variation among different types of primary producers[36]. However, Lys has long retention times in gas chromatography and can co-elute with tyrosine (Tyr), making accurate measurement difficult with typical GC-C-IRMS systems. Similarly, some studies have proposed that the $\delta^{15}N$ difference of Met–Phe can be used to distinguish terrestrial from aquatic ecosystems[37,38]. However, Met tends to co-elute with Glu on polar GC columns and with serine on non-polar columns. Accurate determination of Met $\delta^{15}N$ values therefore requires prior separation of Met from other amino acids by liquid chromatography, followed by derivatization and introduction into the gas chromatograph. Pro is sometimes included in TP calculation formulas, but this practice is not common. As noted above, our measurement system did not allow us to measure Lys, which is paired with Pro, and therefore we did not use TP estimates based on the Pro–Lys pair in our discussion.

#### Validity of the formula to calculate TP

The trophic position (TP) estimated from the nitrogen isotope ratios of Glu and Phe can be expressed as:

$$TP = (\Delta^{15}N_{Glu\text{-}Phe} - \beta)/TDF + 1, \qquad \text{(Equation 1)}$$

where $\Delta^{15}N_{Glu\text{-}Phe}$ is the difference in $\delta^{15}N$ between Glu and Phe; $\beta$ represents $\Delta^{15}N_{Glu\text{-}Phe}$ in primary producers and serves as an offset term that accounts for food web-specific differences; and TDF (trophic discrimination factor) denotes the typical increase in $\Delta^{15}N_{Glu\text{-}Phe}$ associated with a one-step increase in trophic level. This formulation was introduced by Chikaraishi et al.[26]

and has since been widely adopted and empirically validated in numerous studies (reviewed in ref. [27]). In the original formulation[26], TDF is fixed at 7.6. By contrast, $\beta$ is allowed to vary systematically according to the habitat and photosynthetic pathway of the primary producers. Laboratory experiments yielded $\beta$ values of +3.4 for food chains based on aquatic cyanobacteria and algae, −8.4 for those based on terrestrial C3 plants, and +0.4 for those based on terrestrial C4 plants[26]. More recent studies, however, have shown that the principal determinant of $\beta$ is not the photosynthetic pathway (i.e. C3 *vs.* C4) or habitat (i.e. terrestrial *vs.* aquatic), but rather the degree of development of lignin-rich vascular tissues[39,40]. A meta-analysis of stable nitrogen isotopes in terrestrial primary producers further demonstrated substantial overlap in $\Delta^{15}N_{Glu\text{-}Phe}$ (i.e. $\beta$ in Equation 1) between C3 and C4 vascular plants[36]. In other words, in food webs based on vascular C3 or C4 plants, $\beta$ in Equation 1 does not differ in any practically systematic way, and it has become standard practice to use a single $\beta$ value, the one originally defined for C3 plants, to ensure consistency in TP calculations. Indeed, consumers of terrestrial C4 plants yield consistent TP estimates when the conventional $\beta$ for C3 plant ecosystems is applied[27,29]. This conclusion is further supported by CSIA studies of ancient large mammals from archaeological contexts (e.g. refs. [34,41]). Based on these observations, a single $\beta$ value was used, and −8.4 from the original equation proposed by Chikaraishi et al.[26] was used for Equation 1 in this study.

**Precision of CSIA and TP estimates**

For three independent collagen extractants obtained from F051471 (*Cervus*), amino acid derivatization and CSIA for nitrogen were performed to evaluate the precision of the CSIA measurements. For six amino acids, including Glu and Phe, the standard deviation of $\delta^{15}N$ ranged from ±0.5‰ to ±1.2‰, and was ±0.7‰ for both Glu and Phe (**Supplementary Table 6_1**). Repeated measurements of an amino acid standard mixture (**Supplementary Table 6_2**) yielded analytical precision of ±0.3‰ to ±0.9‰ in 1 SD, falling within the typical uncertainty of around ±1‰ reported for CSIA. The standard deviations for the difference between glutamic acid and phenylalanine ($\Delta^{15}N_{Glu\text{–}Phe}$) and for the calculated TP were ±0.2‰ and ±0.0, respectively (**Table 2**), indicating high precision for TP calculation. The calculated TP value of 2.1 for F051471 was consistent with the ecology of *Cervus*, a terrestrial herbivore. These results demonstrate that collagen extraction, amino acid derivatization, and mass spectrometric analysis for CSIA were performed reliably, and that the TP values calculated from $\Delta^{15}N_{Glu\text{–}Phe}$ are precise.

The increase in the $\delta^{15}N$ values of trophic amino acids such as Glu from diet to consumer is represented as the trophic discrimination factor (TDF). Feeding experiments have shown that TDF decreases as the protein content of the diet increases[42]. Compared with the vascular plants consumed by terrestrial herbivores (TP = 2), the animal body tissues eaten by carnivores (TP ≥ 3) generally have higher protein contents, resulting in the potential underestimation of TP for higher terrestrial carnivores. However, feeding experiments on terrestrial animals with diverse dietary compositions indicated that the uncertainty in TP estimates arising from dietary protein content is mostly limited to about ±0.20[26]. In any case, the estimated TP of 2.9 for Penghu 3, which already indicates a highly carnivorous diet, is unlikely to be an overestimate.

### CSIA results of collagen with suboptimal C/N ratios

Typical exogenous organic contaminants that alter the bulk collagen C/N ratio and stable isotope ratios are humic substances, which are rich in carbon but relatively poor in nitrogen[43]. Unlike bulk collagen stable isotope analysis, CSIA involves purification of individual amino acids, thereby greatly reducing the influence of such contaminants[38]. As a result, even bulk collagen samples with C/N ratios outside the acceptable range, $\delta^{15}N_{Glu}$ and $\delta^{15}N_{Phe}$ often retain their endogenous values, in contrast to bulk $\delta^{15}N_{col}$[31,44]. Collagen preservation is generally poor in Penghu fauna, and only 4 of the 72 specimens (5.6%) met the criteria for acceptable quality of bulk collagen. To mitigate data scarcity, we also applied CSIA to two faunal specimens with suboptimal bulk collagen C/N ratios and used the results as supplementary evidence in our discussion.

In the two faunal specimens with suboptimal bulk collagen C/N ratios, $\delta^{15}N_{Phe}$ values were within the same range as in the other Penghu fossils, and the calculated TPs were consistent with values expected for herbivores (**Supplementary Table 6_1**; **Fig. 4**). The extractant collagen, obtained from F051688 (*Equus*), had a bulk collagen C/N ratio of 4.6, but yielded a $\Delta^{15}N_{Glu–Phe}$ of −3.0‰ and a TP of 1.7. For specimen P201508-12 (large-bodied herbivore), the bulk collagen C/N ratio was 3.8, whereas $\Delta^{15}N_{Glu–Phe}$ was −0.4‰ and TP was 2.1. These values match those of the other three Penghu herbivores (**Table 2**). Even for taxa other than *Bubalus* and *Cervus*, faunal specimens with suboptimal bulk collagen C/N ratios yielded plausible CSIA results. These results underscore the robustness of TP estimates based on CSIA for the Penghu fossils.

### Aquatic resource use not supported by CSIA

A high $\Delta^{15}N_{Glu–Phe}$ in CSIA-based TP estimates can arise for two reasons: a high proportion of animal protein in the diet, or a substantial contribution from aquatic resources. The difference in $\delta^{15}N$ values between trophic and source amino acids in primary producers ($\beta$) is generally higher in aquatic systems (+3.4‰) than in terrestrial systems based on vascular plants (−8.4‰: refs. [26,45]). Consequently, animals in aquatic food webs tend to exhibit higher $\Delta^{15}N_{Glu–Phe}$ values than terrestrial animals at the same TP. If an individual consumes food from both terrestrial and aquatic ecosystems, its TP, when calculated using an equation for terrestrial ecosystems, can be overestimated[30,34].

On the other hand, this potential bias can be controlled by comparing the $\delta^{15}N$ values of source amino acids, such as Phe. Aquatic animals typically show lower $\delta^{15}N_{Phe}$ than terrestrial herbivores[30,32,33,35,46]. Thus, individuals that supplement terrestrial foods with aquatic resources can often be identified by $\delta^{15}N_{Phe}$ values lower than those of purely terrestrial animals.

A previous CSIA study of animal skeletons from a Mesolithic site in France, for example, found systematically lower $\delta^{15}N_{Phe}$ in aquatic taxa, such as freshwater turtles and otters, than in terrestrial animals[30]. In the same context, the TP of *Homo sapiens* was calculated as 3.5 ± 0.1 using the TP equation for terrestrial ecosystems, exceeding the theoretical value for terrestrial carnivores (TP = 3), and their $\delta^{15}N_{Phe}$ values were intermediate between those of aquatic and terrestrial animals. These patterns were interpreted as evidence for substantial use of freshwater resources in addition to terrestrial foods[30]. Other previous CSIA studies have reported comparatively high TP values of 2.9 ± 0.2 and 2.8 ± 0.1 calculated for *H. sapiens* from Neolithic France[35] and Bronze–Iron Age Kazakhstan[41], respectively. In those cases, however, their $\delta^{15}N_{Phe}$ values overlapped with those of terrestrial fauna and differed from those of freshwater fish, supporting the conclusion that these populations depended primarily on terrestrial resources.

Even taking these points into account, the CSIA results for Penghu 3 can be parsimoniously explained by strong faunivory on terrestrial herbivores alone, without requiring any contribution from aquatic resources. The TP (2.9) of Penghu 3 derived from the equation for terrestrial ecosystems does not exceed the theoretical value for carnivores (TP = 3) and is therefore not anomalously high (**Fig. 4**). Moreover, the $\delta^{15}N_{Phe}$ value of Penghu 3 falls within the range observed for Penghu terrestrial herbivores (**Fig. 4**).

However, an accurate assessment of any contribution from aquatic resources would require isotopic data from contemporaneous aquatic organisms recovered in the same context. The Penghu fossils were obtained by dredging from the seafloor, and it is inherently difficult to recover the remains of aquatic animals that were actually exploited by hominins. In addition, considering the poor overall preservation of collagen in Penghu fossils, the likelihood of obtaining high-quality collagen from the few available aquatic specimens is low. In any case, combining the CSIA results with the bulk stable isotopic data suggests that aquatic resources did not contribute to the diet of Penghu 3. If aquatic resources contributed to the diet, their contribution must have been too small to be detected isotopically.

**Supplementary Table 6_1. Measured $\delta^{15}N$ values (‰) of individual amino acids and calculated trophic position (TP).**

| NMNS ID | Taxon | Skeletal element | $\delta^{15}N_{Pro}$ | $\delta^{15}N_{Asp+Thr}$ | $\delta^{15}N_{Ser}$ | $\delta^{15}N_{Glu}$ | $\delta^{15}N_{Phe}$ | $\delta^{15}N_{Hyp}$ | $\Delta^{15}N_{Glu-Phe}$ | TP |
|---|---|---|---|---|---|---|---|---|---|---|
| F051713 (Penghu 3) | *Homo* | Tibia | 17.1 | 6.5 | 9.3 | 15.4 | 9.6 | 14.8 | 5.8 | 2.9 |
| F027088 | *Bubalus* | Tibia | 7.7 | 2.1 | 1.4 | 5.8 | 8.2 | 5.2 | -2.4 | 1.8 |
| F027093 | *Bubalus* | Humerus | 14.3 | 8.5 | 6.0 | 11.0 | 12.5 | 6.8 | -1.5 | 1.9 |
| F051471 | *Cervus* | Mandible | 11.3 | 6.1 | 3.5 | 10.0 | 10.1 | 7.6 | -0.1 | 2.1 |
| | | | 12.0 | 6.9 | 5.9 | 11.3 | 11.5 | 8.3 | -0.2 | 2.1 |
| | | | 10.2 | 5.3 | 5.3 | 10.2 | 10.7 | 7.3 | -0.5 | 2.0 |
| F051688 | *Equus* | Tibia or femur | 13.2 | 4.9 | 3.3 | 9.1 | 12.1 | 6.0 | -3.0 | 1.7 |
| P201508-12 | *Palaeoloxodon*? | Limb bone | 16.0 | 4.3 | 2.8 | 9.7 | 10.1 | 5.6 | -0.4 | 2.1 |

**Supplementary Table 6_2. Details of the working amino acid standards used for CSIA.**

| Compound | PN# | $\delta^{13}C$ (‰) | $\delta^{15}N$ (‰) |
|---|---|---|---|
| L-Alanine | AZ104-01 | -17.93 | 43.25 |
| Glycine | AZ6Z2 | -60.02 | -26.63 |
| L-(+)-Norleucine | AZ201 | -28.85 | 18.96 |
| L-Aspartic Acid | AZ203 | -23.95 | 35.2 |
| L-Methionine | AZ2Z0 | -29.4 | -3.79 |
| L-Hydroxyproline | AZ1Z0 | -12.66 | -9.17 |
| L-Leucine | AZ200 | -28.36 | 6.22 |
| L-Glutamic acid | AZ104 | -13.91 | 45.66 |
| L-Phenylalanine | AZ100-01 | -11.2 | 1.7 |

### Supplementary references


1. Tan, K. On the fossil elephant remains in the Government Museum of Taiwan. *Transactions of the Natural History Society of Formosa* **25**, 311-314 (1931).
2. Shikama, T., Otsuka, H. & Tomida, Y. Fossil Proboscidea from Taiwan. *Science Reports of the Yokohama National University, Section II* **22**, 13-62 (1975).
3. Gao, J.-W. Penghu fauna. *Journal of Marine Science (Geology Journal, Institute of Marine Science and Department of Marine Science, Chinese Culture University)* **27**, 123-132 (1982).
4. Chang, C. H. The first fossil record of a short-finned pilot whale (Globicephala macrorhynchus) from the Penghu Channel. *Bulletin of National Museum of Natural Science* **8**, 73-80 (1996).
5. Ho, C. K., Qi, G. Q. & Chang, C. H. A preliminary study of Late Pleistocene carnivore fossils from the Penghu Channel, Taiwan. *Annual of Taiwan Museum* **40**, 1095-1224 (1997).
6. Tsai, C.-H., Fordyce, R. E., Chang, C.-H. & Lin, L.-K. Quaternary Fossil Gray Whales from Taiwan. *Paleontological Research* **18**, 82-93 (2014).
7. Asahara, M., Chang, C.-H., Kimura, J., Son, N. T. & Takai, M. Re-examination of the fossil raccoon dog (Nyctereutes procyonoides) from the Penghu channel, Taiwan, and an age estimation of the Penghu fauna. *Anthropological Science* **123**, 177-184 (2015).
8. Liaw, Y.-L. & Tsai, C.-H. The first fossil of the extant leatherback sea turtle Dermochelys coriacea. *Acta Palaeontologica Polonica* **71**, 267-271 (2026).
9. Biswas, D. S., Banerjee, Y., Baker, D., Chang, C. H. & Tsai, C. H. A glimpse into a vanished ecosystem: reconstructing diet and palaeoenvironment of Palaeoloxodon from the Pleistocene of Taiwan. *R Soc Open Sci* **12**, 250935 (2025).
10. Chang, C. H. *et al.* The first archaic Homo from Taiwan. *Nat. Commun.* **6**, 6037 (2015).
11. Grün, R. & Stringer, C. Direct dating of human fossils and the ever-changing story of human evolution. *Quaternary Science Reviews* **322** (2023).
12. Tseng, Z. J. & Chang, C.-H. A study of new material of Crocuta crocuta ultima (Carnivora: Hyaenidae) from the Quaternary of Taiwan. *Collection and Research* **20**, 9-19 (2007).
13. Tsai, C. H. & Chang, C. H. A right whale (Mysticeti, Balaenidae) from the Pleistocene of Taiwan. *Zoological Lett* **5**, 37 (2019).
14. Tsutaya, T. *et al.* A male Denisovan mandible from Pleistocene Taiwan. *Science* **388**, 176-180 (2025).
15. Kaifu, Y. *et al.* Denisovan leg bones from Taiwan reveal large body size. *bioRxib* (2026).
16. Brock, F. *et al.* Reliability of Nitrogen Content (%N) and Carbon:Nitrogen Atomic Ratios (C:N) as Indicators of Collagen Preservation Suitable for Radiocarbon Dating. *Radiocarbon* **54**, 879-886 (2012).
17. van Klinken, G. J. Bone Collagen Quality Indicators for Palaeodietary and Radiocarbon Measurements. *Journal of Archaeological Science* **26**, 687-695 (1999).
18. DeNiro, M. J. Postmortem preservation and alteration of in vivo bone collagen isotope ratios in relation to palaeodietary reconstruction. *Nature* **317**, 806-809 (1985).
19. Higham, T. *et al.* The earliest evidence for anatomically modern humans in northwestern Europe. *Nature* **479**, 521-524 (2011).

20 Wacker, L., Bollhalder, S., Sookdeo, A. & Synal, H. A. Re-evaluation of the New Oxalic Acid standard with AMS. *Nuclear Instruments and Methods in Physics Research Section B: Beam Interactions with Materials and Atoms* **455**, 178-180 (2019).

21 International Atomic Energy Agency [IAEA] Reference sheet for quality control materials. RS_IAEA-C1 to IAEA-C9.Rev.01 / 2014-03-24. (2014).

22 van der Merwe, N. J. & Vogel, J. C. 13C content of human collagen as a measure of prehistoric diet in woodland North America. *Nature* **276**, 815-816 (1978).

23 Yoneda, M. *et al.* Isotopic evidence of inland-water fishing by a Jomon population excavated from the Boji site, Nagano, Japan. *Journal of Archaeological Science* **31**, 97-107 (2004).

24 Hu, Y. *et al.* Stable isotope dietary analysis of the Tianyuan 1 early modern human. *Proc Natl Acad Sci U S A* **106**, 10971-10974 (2009).

25 Bocherens, H. & Drucker, D. Trophic level isotopic enrichment of carbon and nitrogen in bone collagen: case studies from recent and ancient terrestrial ecosystems. *International Journal of Osteoarchaeology* **13**, 46-53 (2003).

26 Chikaraishi, Y., Ogawa, N. O. & Ohkouchi, N. in *Earth, life, and isotopes* (eds N. Ohkouch, I. Tayasu, & K. Koba) 37–51 (Kyoto University Press, 2010).

27 Ohkouchi, N. *et al.* Advances in the application of amino acid nitrogen isotopic analysis in ecological and biogeochemical studies. *Organic Geochemistry* **113**, 150-174 (2017).

28 McClelland, J. W. & Montoya, J. P. Trophic Relationships and the Nitrogen Isotopic Composition of Amino Acids in Plankton. *Ecology* **83** (2002).

29 Larsen, T., Fernandes, R., Wang, Y. V. & Roberts, P. Reconstructing Hominin Diets with Stable Isotope Analysis of Amino Acids: New Perspectives and Future Directions. *Bioscience* **72**, 618-637 (2022).

30 Naito, Y. I., Chikaraishi, Y., Ohkouchi, N., Drucker, D. G. & Bocherens, H. Nitrogen isotopic composition of collagen amino acids as an indicator of aquatic resource consumption: insights from Mesolithic and Epipalaeolithic archaeological sites in France. *World Archaeology* **45**, 338-359 (2013).

31 Naito, Y. I., Chikaraishi, Y., Ohkouchi, N. & Yoneda, M. Evaluation of carnivory in inland Jomon hunter–gatherers based on nitrogen isotopic compositions of individual amino acids in bone collagen. *Journal of Archaeological Science* **40**, 2913-2923 (2013).

32 Naito, Y. I. *et al.* Ecological niche of Neanderthals from Spy Cave revealed by nitrogen isotopes of individual amino acids in collagen. *J Hum Evol* **93**, 82-90 (2016).

33 Itahashi, Y. *et al.* Amino acid (15) N analysis reveals change in the importance of freshwater resources between the hunter-gatherer and farmer in the Neolithic upper Tigris. *Am J Phys Anthropol* **168**, 676-686 (2019).

34 Tejada, J. V. *et al.* Isotope data from amino acids indicate Darwin's ground sloth was not an herbivore. *Scientific reports* **11**, 18944 (2021).

35 Rey, L. *et al.* Specifying subsistence strategies of early farmers: New results from compound-specific isotopic analysis of amino acids. *International Journal of Osteoarchaeology* **32**, 654-668 (2022).

36 Ramirez, M. D., Besser, A. C., Newsome, S. D. & McMahon, K. W. Meta-analysis of primary producer amino acid δ15N values and their influence on trophic position estimation. *Methods in Ecology and Evolution* **12**, 1750-1767 (2021).

37 Ishikawa, N. F. *et al.* A new analytical method for determination of the nitrogen isotopic composition of methionine: Its application to aquatic ecosystems with mixed resources. *Limnology and Oceanography: Methods* **16**, 607-620 (2018).

38 Ohkouchi, N. A new era of isotope ecology: Nitrogen isotope ratio of amino acids as an approach for unraveling modern and ancient food web. *Proc Jpn Acad Ser B Phys Biol Sci* **99**, 131-154 (2023).

39 Ohkouchi, N. & Takano, Y. in *Treatise on Geochemistry* (eds. H. D. Holland & K. K. Turekian) 251-289 (Elsevier Science, 2014).

40 Kendall, I. P. *et al.* Compound-specific delta(15)N values express differences in amino acid metabolism in plants of varying lignin content. *Phytochemistry* **161**, 130-138 (2019).

41 Itahashi, Y. *et al.* Dietary diversity of Bronze-Iron Age populations of Kazakhstan quantitatively estimated through the compound-specific nitrogen analysis of amino acids. *Journal of Archaeological Science: Reports* **33** (2020).

42 Whiteman, J. P., Rodriguez Curras, M., Feeser, K. L. & Newsome, S. D. Dietary protein content and digestibility influences discrimination of amino acid nitrogen isotope values in a terrestrial omnivorous mammal. *Rapid Commun Mass Spectrom* **35**, e9073 (2021).

43 Schwarcz, H. P. & Nahal, H. Theoretical and observed C/N ratios in human bone collagen. *Journal of Archaeological Science* **131** (2021).

44 Itahashi, Y. *et al.* Preference for fish in a Neolithic hunter-gatherer community of the upper Tigris, elucidated by amino acid δ15N analysis. *Journal of Archaeological Science* **82**, 40-49 (2017).

45 Naito, Y. I. *et al.* An overview of methods used for the detection of aquatic resource consumption by humans: Compound-specific delta N-15 analysis of amino acids in archaeological materials. *Journal of Archaeological Science: Reports* **6**, 720-732 (2016).

46 Fontanals-Coll, M. *et al.* Stable isotope analyses of amino acids reveal the importance of aquatic resources to Mediterranean coastal hunter-gatherers. *Proc Biol Sci* **290**, 20221330 (2023).
